# Genomic Signatures of Microgeographic Adaptation in *Anopheles coluzzii* Along an Anthropogenic Gradient in Gabon

**DOI:** 10.1101/2024.05.16.594472

**Authors:** Josquin Daron, Lemonde Bouafou, Jacob A. Tennessen, Nil Rahola, Boris Makanga, Ousman Akone-Ella, Marc F Ngangue, Neil M. Longo Pendy, Christophe Paupy, Daniel E. Neafsey, Michael C. Fontaine, Diego Ayala

## Abstract

Species distributed across heterogeneous environments often evolve locally adapted populations, but understanding how these persist in the presence of homogenizing gene flow remains puzzling. In Gabon, *Anopheles coluzzii,* a major African malaria mosquito is found along an ecological gradient, including a sylvatic population, away of any human presence. This study identifies into the genomic signatures of local adaptation in populations from distinct environments including the urban area of Libreville, and two proximate sites 10km apart in the La Lopé National Park (LLP), a village and its sylvatic neighborhood. Whole genome re-sequencing of 96 mosquitoes unveiled *∼*5.7millions high-quality single nucleotide polymorphisms. Coalescent-based demographic analyses suggest an *∼*8,000-year-old divergence between Libreville and La Lopé populations, followed by a secondary contact (*∼*4,000 ybp) resulting in asymmetric effective gene flow. The urban population displayed reduced effective size, evidence of inbreeding, and strong selection pressures for adaptation to urban settings, as suggested by the hard selective sweeps associated with genes involved in detoxification and insecticide resistance. In contrast, the two geographically proximate LLP populations showed larger effective sizes, and distinctive genomic differences in selective signals, notably soft-selective sweeps on the standing genetic variation. Although neutral loci and chromosomal inversions failed to discriminate between LLP populations, our findings support that microgeographic adaptation can swiftly emerge through selection on standing genetic variation despite high gene flow. This study contributes to the growing understanding of evolution of populations in heterogeneous environments amid ongoing gene flow and how major malaria mosquitoes adapt to human.

**Significance:** *Anopheles coluzzii*, a major African malaria vector, thrives from humid rainforests to dry savannahs and coastal areas. This ecological success is linked to its close association with domestic settings, with human playing significant roles in driving the recent urban evolution of this mosquito. Our research explores the assumption that these mosquitoes are strictly dependent on human habitats, by conducting whole-genome sequencing on *An. coluzzii* specimens from urban, rural, and sylvatic sites in Gabon. We found that urban mosquitoes show *de novo* genetic signatures of human-driven vector control, while rural and sylvatic mosquitoes exhibit distinctive genetic evidence of local adaptations derived from standing genetic variation. Understanding adaptation mechanisms of this mosquito is therefore crucial to predict evolution of vector control strategies.

## Introduction

The genetic mechanisms by which natural populations can adapt to heterogeneous habitats has been a fundamental question in ecology and evolution. Local adaptation is responsible for the acquisition of traits providing a selective advantage under specific environmental conditions, regardless of the fitness consequences in other habitats (Kawecki and Ebert 2004; Tiffin and Ross-Ibarra 2014). At the molecular level, local adaptation can be associated with *de novo* mutation where advantageous alleles can quickly reach fixation through “hard” selective sweeps. However, this process can be relatively slow to evolve. Instead, local adaptation can emerge from the recycling of the preexisting standing genetic variants (SGV) through “soft” selective sweeps (Louis et al. 2021; Small et al. 2023). Because hard selective sweeps of *de novo* mutations are easier to detect using outlier approaches (Smith and Haigh 1974), the vast majority of the empirical work has been centered upon a “hard-sweep” model of adaptation. However, a growing literature suggests that soft sweeps might be a more frequent mode of adaptation in many natural populations (Hermisson and Pennings 2005; Messer and Petrov 2013; Garud et al. 2015; Sheehan and Song 2016; Schrider and Kern 2017). More importantly, both selection modes (hard and soft) are fully compatible and complementary. Consequently, understanding the relative interplay between new mutations and SGV for the adaption process is a key challenge to understand the ability of species to evolve into changing environments (Visser 2008).

The African malaria mosquito *Anopheles coluzzii* exhibits a remarkable ecological ability to proliferate in a broad range of habitats as diverse as humid rain-forest, highland and dry savannah (Simard et al. 2009; Tene Fossog et al. 2015). The ecological success of *An. coluzzii* is directly rooted by the tremendous genetic and chromosomal polymorphisms and in its close association with human settings (Ayala et al. 2014; Fontaine et al. 2015a; Ayala et al. 2017; Anopheles gambiae 1000 Genomes Consortium et al. 2020). Humans provide blood meals, as well as shelters, and breeding sites, to *An. coluzzii* (White et al. 2011). This degree of specialization on humans has drastically impacted the recent evolution of *An. coluzzii* (Ayala and Coluzzi 2005). First, the adaptation to newly modified landscape by the advent of agriculture 5,000 - 10,000 years ago (ya) and human density impacted its evolution by greatly increasing their population sizes (Ag1000G Consortium 2017; Ag1000G Consortium et al. 2020). Second, the massive use of insecticides for malaria control in the second half of the last century selected resistant variants within natural populations of this mosquito by genetic changes at targeted sites in the genome (i.e., *de novo* mutations), metabolic, and behavioral modifications (i.e. biting rhythm (Sangbakembi-Ngounou et al. 2022). Third, the rapid and recent development of large cities in Africa led to strong selection of genetic and physiological factors (i.e. osmoregulation (Tene Fossog et al. 2015)) to tolerate urban pollution through detoxification (Kamdem et al. 2017). Therefore, humans are key actors on the environmental adaptation of this mosquito.

Motivated by the profound implications for human health, malaria research has intentionally narrowed its focus to anthropogenic settings, neglecting less anthropized and wild areas (Ayala et al. 2009; Kyalo et al. 2017). In the last few years, several studies incidentally reported the presence of species within the *An. gambiae* complex in wild conditions away from any kind of permanent human presence in Gabon, South Africa or Madagascar (Paupy et al. 2013; Munhenga et al. 2014; Zohdy et al. 2016), but also even in well studied areas such as in Burkina-Faso (Tennessen et al. 2021). For example, the discovery of a previously unknown species within the *An. gambiae* complex*, An. fontenillei*, but present away from anthropic settings, underscores our incomplete understanding of biodiversity in *Anopheles* (Barrón et al. 2019). Although unnoticed, these observations challenge the assumption that these mosquitoes are strictly bound to anthropogenic habitats. These observations also raise questions about the evolutionary processes involved in the local adaptation of these mosquitoes in the absence of their main hosts, humans, supporting their extraordinary ability to adapt to a large variety of eco-anthropogenic conditions. Therefore, wild areas, such as National Parks or protected areas, provide a compelling opportunity to investigate the origin and evolution of the main malaria mosquitoes in Africa and possibly also about factors underlying their vectorial capacity.

Here, we report an in-depth characterization of the genetic variation of *An. coluzzii* along an anthropogenic gradient in Gabon, Central Africa. By comparing sylvatic and rural populations in the La Lopé National Park (LLP), together with an urban population from Libreville (LBV) and also, at a broader scale in Africa, with the data from the *An. gambiae* 1000 genome (Ag1000G) project, we investigated the evolutionary processes by which these mosquitoes adapted to these contrasted environments. Given the permanent and stable occurrence of *An. coluzzii* not only in the rural villages of LLP, but also in the proximate forested areas (Ayala et al. 2009; Paupy et al. 2013; Barrón et al. 2019), we hypothesized that several genetic determinants may be involved in contrasted adaptations to these environments (Figure 1A). To address these questions, we sequenced the whole genome of 96 mosquitoes sampled across the anthropogenic gradient in Gabon. We investigated the population genetic variation, its spatial structure, and the interplay between evolutionary forces (genetic drift, migration and selection) shaping the population genetic variation along this gradient. We identified very contrasted dynamics between the urban LBV population and the populations from LLP. On one side, the urban population exhibited clear evidence of strong selective pressure marked by hard selective sweeps associated with adaptation to polluted waters and insecticide, typical of a highly populated and anthropized environment. On the other side, the populations from the LLP exhibited much more diffuse genetic evidence of selection on SGV. Despite the close proximity (∼10km) of the rural and sylvatic populations, we found these two populations shared the same neutral genetic background, yet displaying selective evidence of local adaptation to the contrasted selective pressures of each environment. Together this study provides a unique opportunity to evaluate the action of local adaptation in the face of a strong gene flow on a genomic scale. Our results suggest that the exceptional SGV of this species is key to adaptation to sylvatic conditions.

**Figure 1:**
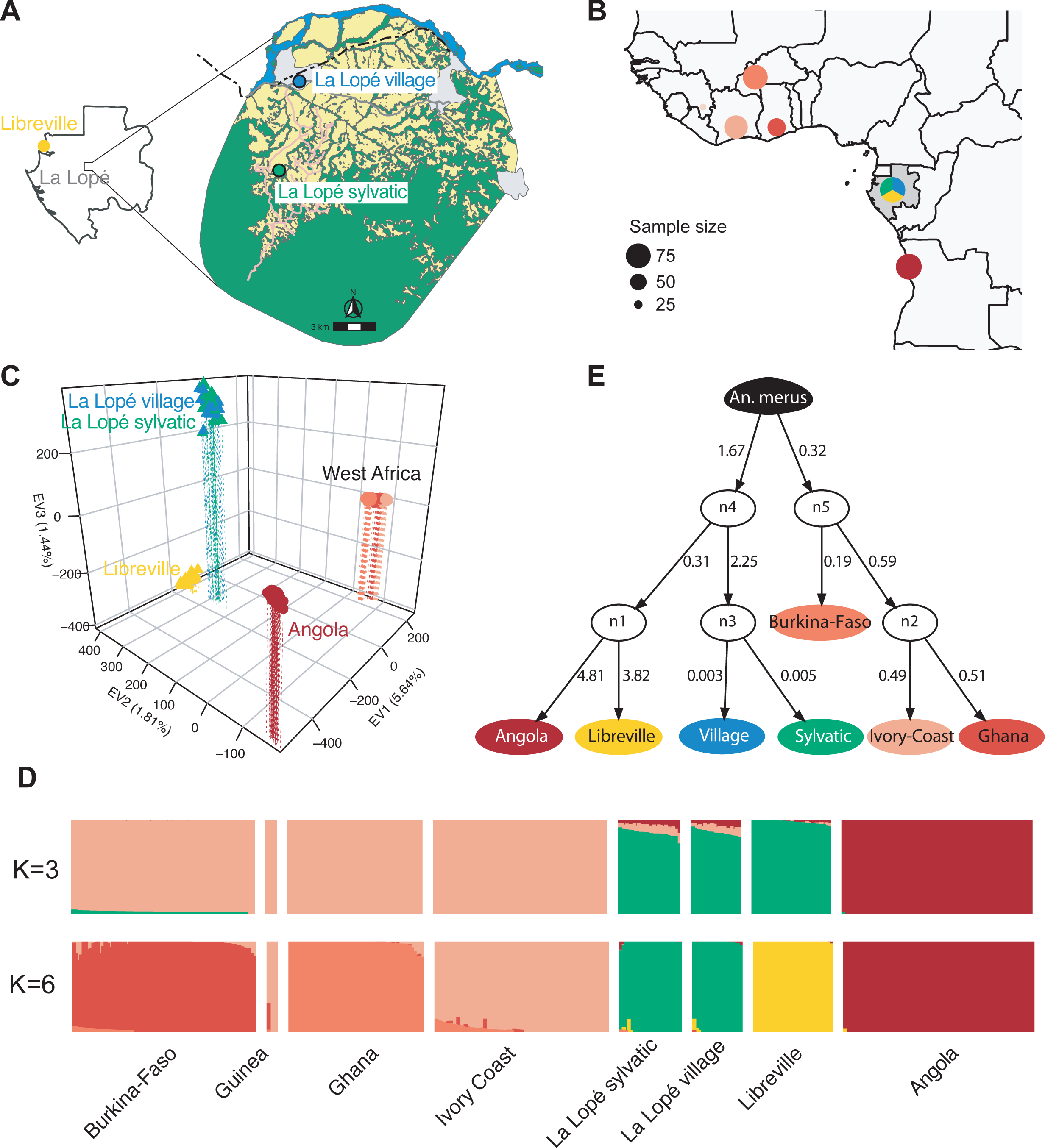
Geographic sampling and population structure and admixture. **(A)** Sampling locations of the three focal populations in Gabon. A total of 96 *An. coluzzii* mosquitoes were collected in Libreville as larvae, or as adults in the village and sylvatic area of the La Lopé National Park (see Supplementary Table S1 for details). The geographic map of the National Park of La Lopé highlights the contrasted habitats that coexist in the park, with stable and permanent settlement of mosquitos observed in the village and the sylvatic areas. The base map was produced by digitizing the Gabonese land use map freely available from the *Agence Nationale d’Étude et d’Observation Spatiale du Gabon* (http://ageos.ga). **(B)** Geographic locations of the 283 *An. coluzzii* mosquitoes analyzed for the population structure at the scale of West Africa. Colors of regional groups are consistent throughout the study. The size of sampling point is proportional to the sample size. **(C)** Population genetic structure captured by the top three principal components of the PCA (see also the scree-plot on Supplementary Figure 2A). **(D)** *ADMIXTURE* analyses of the *An. coluzzii* populations. Each bar shows the genetic ancestry proportions for each individual to each genetic cluster tested. The best fitting model according to the cross-validation error rates (Supplementary Figure S5B) was observed at K=3, but finer sub-structuration was clearly visible until K=6. See Supplementary Figure 3 for the different solution from K=2 to 9, and the associated cross-validation (CV) error-rates. **(E)** Most likely population graph topology recovered with a posterior probability of 95.04% using *AdmixtureBayes* (Nielsen et al. 2023). The population graph was rooted using *An. merus* used as outgroup species. The graph is composed of leaf nodes (supported with posterior probabilities higher than 0.98) that are not the product of an admixture event (white circles). The numbers on the branch connecting populations capture the amount of genetic drift between populations.

## Results

### Highly shared sanding genetic variation between sylvatic, rural and urban populations in Gabon

We analyzed the whole genome variation at single-nucleotide polymorphisms (SNPs) by resequencing 96 *An. coluzzii* mosquito samples coming from three locations in Gabon (n=32 in each) along the anthropogenic gradient: the urban area of Libreville city (LBV) and two locations in the La Lopé National Park including a rural village area (LPV) and a sylvatic area 10km to the South (LPS) (Figure 1A). After read cleaning, mapping, SNP variant calling, and individual quality filtering, we retained a total of 87 individuals with an average sequencing depth coverage of 53.1 ± 25.01 (see the methods and Supplementary Figure S1, Figure S2, and Table S1). Nine additional individuals were removed because they exhibited a missingness rate over 10% (after genotype quality filtering) as well as one extra individual due to a high degree of relatedness (Supplementary Figure S3 and Table S1). The final dataset thus included 77 individuals, sequenced at a depth coverage of 57.1 (52.4) ± 23.2 [23.1 – 141.9], including 61 females and 16 males (Table S1). A total of 5.9 million high-quality SNPs were obtained after data quality check and filtering. The SNP density along the genome included 1 SNP every 25 bp on average, with most of the SNPs being shared among samples from the three geographic sites (Supplementary Figure S2C). This indicates a high level of shared standing genetic variation (SGV).

### Strong genetic differentiation between mosquitoes from Libreville and La Lopé, contrasting with uniform genetic profiles across La Lopé rural and sylvatic

Next, we investigated the genetic structure of the Gabonese samples alone, and also in the context of the continent-wide population structure of *An. coluzzii* by combining our data set with those from *An. coluzzii* of the phase-2 *Ag1000G* consortium project (Ag1000G Consortium et al. 2020) (Figure 1B). Focusing only on the Gabonese samples, the principal component analysis (PCA) showed that only one PC axis was meaningful and revealed that the urban mosquitoes from Libreville (LBV) grouped tightly together and apart from those of the La Lopé National Park (Supplementary Figure 4B). Considering them with the other *An. coluzzii* populations from the Ag1000G, the PCA showed that the top three PC axes captured a disproportionate amount (∼9%) of the total variance compared to the remaining PC axes (Supplementary Figure 4A). The individual PC scores (Figure 1C) revealed that the Gabonese populations were well distinct from those of Angola and from those of North-Western Africa (Burkina-Faso, Ghana, Ivory Coast, Guinea) in the plan of the two first PC axes. The third axis further split the urban LBV mosquitoes from those of LLP. In contrast, the two proximate LLP populations could not be discriminated (Figure 1C), at least based on the neutral unlinked SNPs used to study the genetic structure (Supplementary Figure S3B). The genetic ancestry analysis of ADMIXTURE revealed a similar genetic structure as the PCA. Three main genetic clusters (K=3) were identified as best-fitting solution, which minimized the cross-validation error rates (Figure 1D, Supplementary Figure S5). These three genetic clusters were composed of *An. coluzzii* mosquitoes from Gabon, Angola, and NW Africa. However, additional sub-structure was clearly visible at higher K values and consistent with the PCA and previous results from the (Ag1000G Consortium et al. 2020). In fact, up to six distinct genetic groups were identified including Angola, LBV, and the two LLP populations identified as a single genetic cluster, Burkina-Faso, Ghana, and Ivory Coast and Guinea; the latter two also shared the same ancestry.

Genetic differentiation expressed as *F_ST_* values among population pairs were all significantly different from zero, but one: the sylvatic and village populations of the LLP National Park (p=0.575; Supplementary Figure S6). Globally, *F_ST_* values recovered similar genetic structuration as captured by the PCA and the ADMIXTURE analyses. Aside from the Guinean population with a very small sample size (n=5), all populations from NW Africa displayed shallow but significant genetic differentiation with *F_ST_* values ≤ 0.01 (*p* ≤ 0.05). The highest *F_ST_* values were observed when comparing mosquitoes from Angola and those from NW Africa (*F_ST_* > 0.13; *p* ≤ 1.25e^-17^). Comparisons involving the Gabonese populations (LBV and LLP) displayed intermediate *F_ST_* values (0.05) (Supplementary Figure S6).

We explored further the genetic ancestry relationships among the different populations of *An. coluzzii* and determined whether they may descend or not from admixture event(s) using *AdmixtureBayses* (Nielsen et al. 2023). The best-fitting population graph (Figure 1E) collected 95% of the posterior probability and exhibited no admixture event among populations of *An. coluzzii*. All the nodes on that graph were highly supported with posterior probabilities >98%. The graph suggested that the populations from NW African localities (Burkina-Faso, Ivory Coast and Ghana) were all closely related to each other, as shown by the low estimates of genetic drift values along the branches of the graph and as suggested in the PCA and previous studies (Ag1000G Consortium et al. 2020). Populations from Central Africa (Angola and Gabon) were also more closely to each other than those from NW Africa, as shown by the strong support on the node (node *n4*). Interestingly, *AdmixtureBayses* analysis suggested that populations from LBV and Angola were more closely related to each other than with the two LLP populations (node *n1*). Nevertheless, both Angola and LBV displayed elevated genetic drift values from each other, suggesting important differentiation between them and small effective population sizes (*Ne*). The two LLP localities were very closely related to each other with very low drift values from the ancestral node (<0.005, *n3*). They coalesced deeper in the tree with the ancestor of the (Angola, LBV) ancestor on the node *n4*.

One consistent observation identified by all analyses so far was that samples from the two proximate LLP localities (*i.e.,* village and sylvatic) remained nearly undistinguishable at neutral unlinked SNPs suggesting they belong to a same genetic pool (Figure 1C, Figure 1D, K=6 in Figure 1D and Supplementary Figure S5). We tested formally this hypothesis at neutral loci by comparing the observed joined site-frequency spectrum (jSFS) with a simulated jSFS expected under panmixia (obtained by permuting sample labels, see Materials and Methods) using *δaδi* (Gutenkunst et al. 2009). The observed jSFS did not statistically depart from the null distribution generated by 1000 permuted jSFS (p-value > 0.45; Supplementary Figure S7), confirming the genetic homogeneity observed with the PCA, ADMIXTURE, FST, and *AdmixtureGraph*. A similar analysis performed between populations from Libreville and either of the village and sylvatic LLP areas shows highly significant differences (p-value < 10^-26^ for each test; Supplementary Figure S7). Accordingly, average *F_ST_* values between populations from LBV and either the rural or sylvatic areas of LLP were 0.05, one order of magnitude higher than between the two localities in La Lopé (*F_ST_* = 1.7e-4, Supplementary Figure S6).

### Reduced diversity and increased autozygosity in Gabonese mosquito populations typical of Central African An. coluzzii

We characterized further the genetic diversity of the sampled populations in the three Gabonese localities in comparison with those from the Ag1000G using a variety of summary statistics (Figure 2). Overall, the summary statistics showed that the samples from the three Gabonese localities were very similar to the population from Angola in Central Africa. In contrast with populations from NW Africa, *An. coluzzii* populations from Gabon and Angola displayed lower nucleotide diversity, Tajima’s D values closer to 0, slower LD-decay, low amounts of rare frequency variants in their SFS’s, larger proportions of individual genomes covered by long runs of homozygosity (*F_ROH_* > 100kb), and longer inter-individual identical-by-descent (IBD) tract lengths (Figure 2). All these distinctive patterns of genetic diversity suggest that the population dynamics and history of *An. coluzzii* are very distinct in Central Africa compared to those in NW Africa, with populations of smaller effective population size (*Ne*), with possibly more stable demography with small *Ne* values or strongly fluctuating *Ne* size, also inducing some level of inbreeding (as suggested by the F_ROH (100kb)_ >0.10). Of interest is the LBV population which displayed an even higher inbreeding level compared to the sylvatic and rural populations of LLP as suggested by the higher *F_ROH_* values, slightly higher Tajima’s D values, slower LD-decay, and smaller nucleotide diversity. This is consistent with smaller and possibly more fluctuating population *Ne* value in LBV than the mosquitoes from LLP.

**Figure 2:**
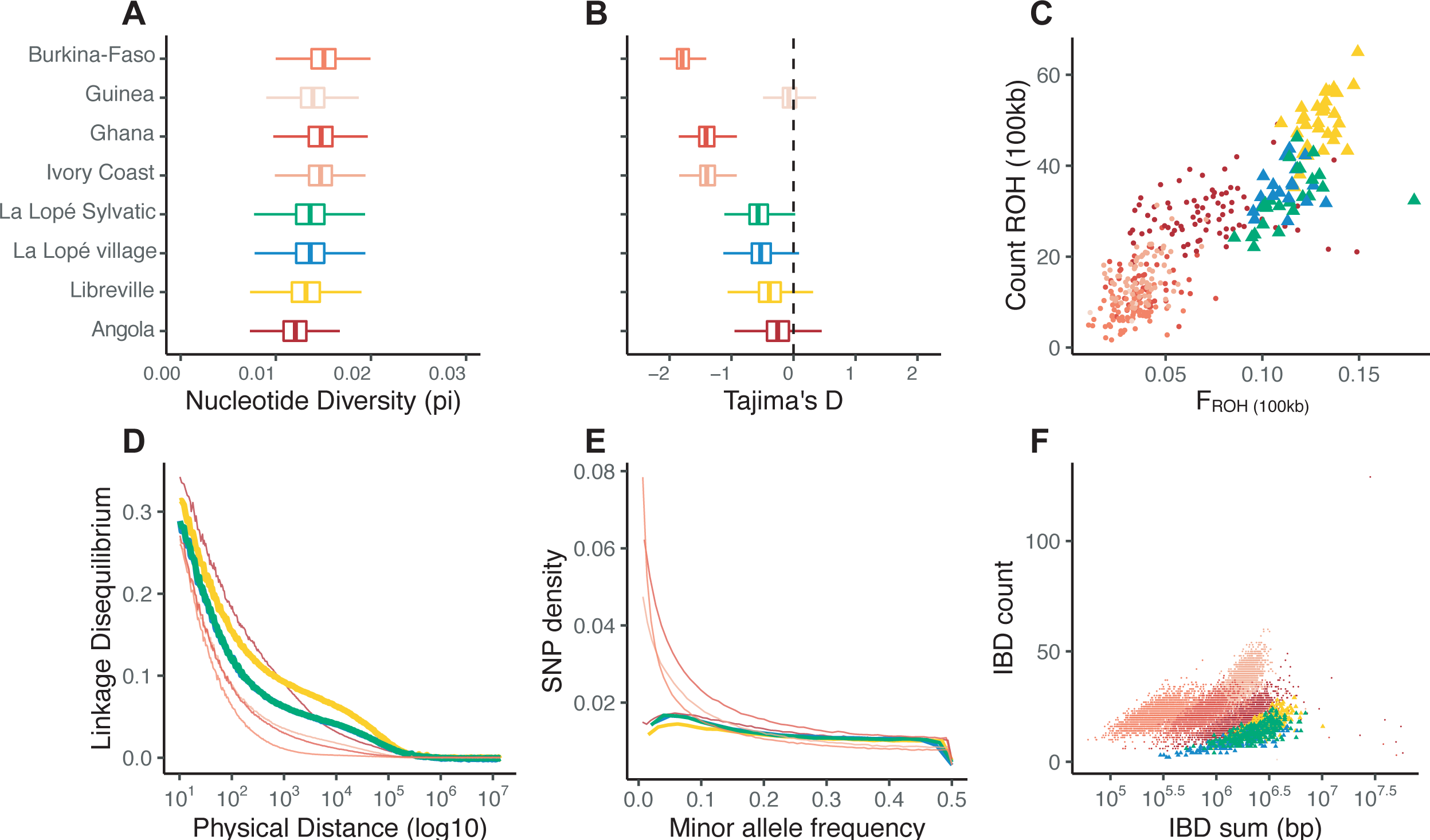
Genetic diversity of the African populations of *An. coluzzii*. Boxplots showing **(A)** the nucleotide diversity (π) and **(B)** the Tajima’s D estimated in 10 kb non-overlapping windows. **(C)** Count and frequency of the runs of homozygosity (ROH) ≥ 100kb observed in the individual mosquitoes. Each point represents an individual mosquito. **(D)** Decay in linkage disequilibrium (*r^2^*) as a function of the physical distance between SNPs. **(E)** Minor allele frequency spectrum (MAF). **(F)** Scatterplot of the count versus the sum of runs of identity by descent (IBD) between individuals, with each dot representing a pair of individuals drawn from the same population.

### Small and relatively stable demographic history of Gabonese populations

We estimated the demographic history of each population of *An. coluzzii* in Gabon, comparatively with those from the *Ag1000G*, first using *Stairway plot* v2 (Liu and Fu 2020) and the unfolded (*a.k.a.* polarized) SFS of the putatively neutral and recombining portion of chromosome 3 (Figure 3A and 3B). The two proximate populations from LLP shared similar historical trajectories in *Ne* variation, with 2 consecutive bottlenecks, one *c.*40k years ago (kya), and another one *c.*20 kya. *Ne* values remained relatively stable between the bottlenecks and since the past thousands years with however consistent evidence of a very recent decline. Noteworthy was the population from LBV, which displayed consistently lower effective sizes compared to the rural and sylvatic LLP populations, in line with the genetic diversity estimates (Figure 2). The effective population sizes of the three Gabonese populations (Figure 3A) were also comparable to those observed in Angola, and were at least one order of magnitude smaller than those from NW Africa (Figure 3B), also consistent with genetic diversity estimates and previous studies (Ag1000G Consortium et al. 2020).

**Figure 3:**
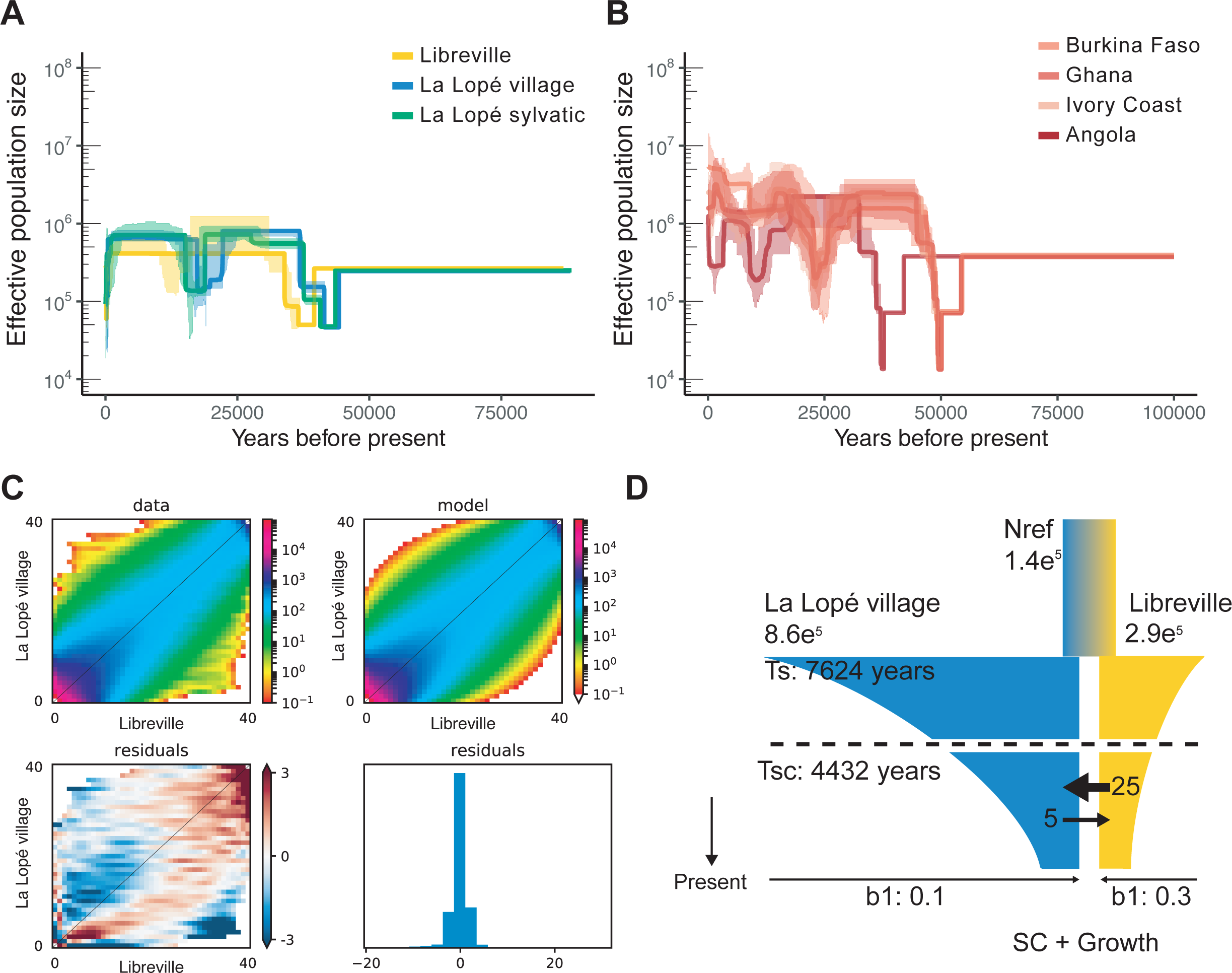
Demographic histories of the African populations of *An. coluzzii* estimated from genetic data. **(A-B)** Historical changes in effective population size (*Ne*) were inferred via *StairwayPlots* 2 (Liu and Fu 2020) for the three Gabonese populations **(A)** and for populations of the Ag1000G consortium (Ag1000G Consortium 2017) from Western Africa and Angola provided here as a comparison **(B)**. The *Ne* values for of each population was rescaled using a generation time (g) of 11 years and a mutation rate (μ) of 3.5 × 10^−8^ per site and per year. The main-colored lines show the median estimates and light shade areas represent 95% confidence intervals. **(C)** Results of *∂a∂i* (Gutenkunst et al. 2009) 2 population analysis. The top panel represents the observed joint site frequency spectrum (jSFS) for Libreville versus La Lopé village along with a secondary contact model fit (top right panel) and distributions of the residuals (bottom panel). **(D)** Visual representation of demographic model diagram fitting the best model to the jSFS of the populations from Libreville *versus* La Lopé village, together with the parameter estimates of the model. (see Supplementary Table S3 for details on parameters estimations)

### Secondary contact and asymmetric gene flow have homogenized Gabonese populations

To better understand the isolation history between the populations from LBV and LLP, we tested which isolation models best fitted the observed unfolded joined SFS (jSFS) using *δaδi* (Gutenkunst et al. 2009). We compared 8 distinct models of population isolation differing in terms of *Ne* variation (constant *vs* exponentially, -G), and in terms of gene flow history contrasting strict isolation without gene-flow (–SI), isolation with continuous migration (–IM), ancestral migration (– AM), or secondary contact (– SC). The best fitting model displaying the lowest log-likelihood scores and best AIC criterion identified the scenario implying a secondary contact with population-size change (SC+G) (Supplementary Figure 8 and Supplementary Tables 2). This demographic scenario suggested that the isolation between LBV and LLP involved a period of allopatric divergence followed by a secondary contact (SC), with a slight asymmetric gene flow predominantly from LBV into LLP, and an exponential decline in both populations (Figure 3C). Model parameter estimates considering this SC+G model (Figure 3D and Supplementary Table 3) suggest that the populations from Libreville city (LBV) and La Lopé (LLP) split *c.* 7,624 years ago (*Ts*) from an ancestral population of 142,332 individuals (*Nref*) into two populations, each one with distinct *Ne* values, respectively of 292,460 and 865,159 individuals for LBV and LLP. The two populations would have remained isolated until *c.* 4,432 years ago (*T_sc_*), time at which a secondary contact would have restored an asymmetric gene flow with five times more effective migrants from LBV into LLP than in the reverse direction (Figure 3D and Supplementary Table 2).

### Contrasted regimes of positive selection along the anthropogenic gradient

We screened the genome for genomic evidence of positive selection that may have contributed to local population adaptation across the sylvatic, rural, and urban anthropogenic gradient. We first conducted genome scan comparing population pairs based on differences in long-range haplotype homozygosity (*XP-EHH* (Sabeti et al. 2007) and differences in allele frequency (*F_ST_* - based statistics) (Alexander et al. 2009). *XP-EHH* detects differences in extended runs of haplotype homozygosity between populations, reflecting variation in haplotype length and LD along the genome (Voight et al. 2006; Sabeti et al. 2007), tailored to detect recent selective sweeps, while *F_ST_* statistics scans can complement haplotype-based scans by identifying outliers displaying significant differentiation in allele frequencies between populations along their genomes. The comparison between LBV and either rural or sylvatic LLP populations revealed a few clear and strong signals of positive selection in the urban group at both *F_ST_* (Supplementary figure S9) and *XP-EHH* (Figure 4A and Supplementary figure S10) statistics. Those selection footprints were found centered mostly around well-known genomic regions harboring insecticide resistance genes, jointly detected by *XP-EHH* and *F_ST_* statistics (*VGSC*, *GABA* and *GSTE*) or based on *F_ST_* genome scan only (cytochrome *P450s cyp6p* and *cyp9k1*). Additionally, away from the known resistance genes, a few other genomic regions exhibited also significant evidence of strong positive selection in the urban LBV population. Among them, one stretch between positions 41,275,000 and 41,500,000 on the chromosome 3L consistently stood out through the comparisons with both LPV and LPS population (Figure 4A). The four annotated genes in this region included the gene *AGAP012385* encoding for a Toll-like receptor signaling pathway known to mediate anti-pathogen defense, including against *Plasmodium* (Clayton et al. 2013). Twenty-six significant SNPs were found in the 5kb upstream and downstream regions of that gene. Positive selection was also clearly suggested by both *XP-EHH* and *F_ST_* statistics in the pericentromeric region of chromosome 2L and 2R. This region is also well known for its selection history associated with the *Voltage-Gated Sodium Channel (VGSC)*, known as ‘*kdr*’ owing to their knock-down-resistance phenotype, reduce susceptibility to DDT and pyrethroid in *Anopheles* (Davies et al. 2007; Ag1000G Consortium 2017; Lynd et al. 2018).

**Figure 4:**
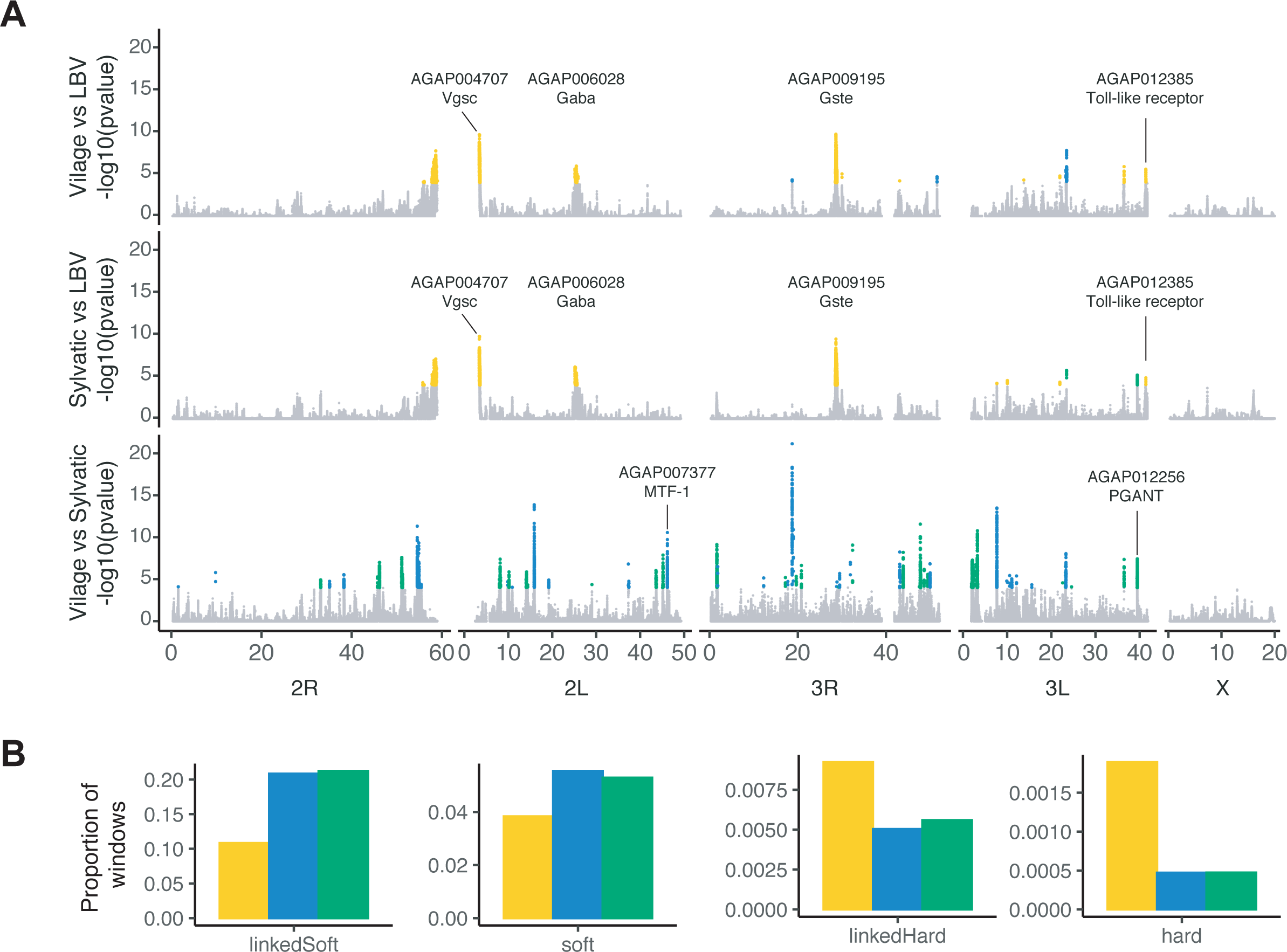
Signals of positive selection among Gabonese populations. **(A)** Genome scans of the XP-EHH statistic *p*-values calculated at the SNP level along the genome and plotted for all pairwise populations comparisons. Each dot is colored to denote its significance, with gray dots indicating non-significant SNPs and colored dots representing significant value (pvalue<1e-4) suggesting positive selection. The color of the dot corresponds to the population in which the SNP has been found significant. **(B)** Proportion of the overall genomic windows (N=12,693) classified within the four different class sweeps in the DiploS/HIC analysis. Color code of the populations is consistent with Figure 1.

By contrast, *XP-EHH* values between the village and sylvatic area of LLP yielded 44 significant outlying regions exhibiting positive signatures of selection scattered across the autosomes (Figure 4 and Supplementary figure S10). The proportion of significant SNPs in each population did not differ significantly between the sylvatic and rural populations of LLP (Supplementary Figure S11, χ2 test, *p-value* > 0.05). Those selective footprints were exclusively detected by the *XP-EHH* statistic, while *F_ST_* scans revealed almost no differentiation in allelic frequency along the entire genome of the sylvatic and rural LLP populations (Supplementary Figure S9). This contrast between *XP-EHH* and *F_ST_* is consistent with very recent selection signals, most likely involving soft selective sweeps on the SGV. The *XP-EHH* test can detect selective sweeps in which the selected allele has risen to high frequency or fixation in one population, but remains polymorphic in the human population as a whole (Sabeti et al. 2007). Under this circumstance, *XP-EHH* can detect signals in regions that do not display any outlying *F_ST_* values. *F_ST_*-based selection scan is meant to detect signals of excessive levels of genetic differentiation, which would typically capture hard selective sweeps at known resistance genes discriminating the city from the rural area. However, in the case of soft sweep and selection on the SGV, differences in allele frequency becomes much more subtle, and the power to detect significant outlying *F_ST_* value drop as well (Voight et al. 2006; Sabeti et al. 2007). Since the populations from the village and sylvatic area of LLP are derived from a same genetic pool and exhibit similar demographic histories and LD-decay, the sensitivity of haplotype-based methods is maximized to identify recent processes of positive selection (Ma et al. 2015; Klassmann and Gautier 2022).

Annotating the SNP effect using *snpeff* (Cingolani 2022), for the significant SNPs identified with *XP-EHH* scans shows that variant sets were significantly different in the nature of their annotation (Supplementary Figure S11A, *χ^2^* test =538.562, *p-value* < 1e^-5^). Selected variants distinguishing the LBV from the rural LLP area were found enriched in synonymous SNPs, while variants distinguishing the village from the sylvatic LLP populations were found enriched at intergenic regions (Supplementary Figure S11B). Together, our results highlight the contrasted nature of the positive selection signal across our pairwise comparison. On one hand, between the urban LBV population and the rural LLP populations, a strong selection signal is detected, extremely localized at very few loci. On the other hand, between the village and the sylvatic LLP populations, the signal is spread out through the entire genome encompassing few SNPs per picks.

We explored further the hypothesis that urban population from LBV and the two proximate localities in the LLP National Park exhibit distinct selective regimes: the first one being dominated by hard-selective sweeps, and the two others by soft-sweep on the SGV. For that purpose, we used the supervised deep learning technique of *DiploS/HIC* (Kern & Schrider, 2018). This approach uses coalescent simulations of genomic regions based on the observed demographic history for each population as training set. From these simulations, this approach uses a deep learning algorithm to classify genomic windows into 5 categories of positive selection (neutral, hard, linked-hard, soft, and linked-soft selective sweeps) based on patterns of genetic variation surrounding a focal genomic region of 110kb subdivided into 11 windows of 10kb. High level of convergence was observed for the summary statistics of our simulated dataset compared to the empirical data under the different selection categories (Supplementary Figure S12). The performance of the Convoluted Neural Network (CNN) classifier to discriminate among the five types of selection categories based on these simulations was also quite good as shown by the confusion matrix (Supplementary Figure S13). In average over the three populations, we estimated respectively a 76% and a 61% accuracy to correctly predict hard and soft selective sweeps. Hard and soft sweeps were found to be clearly distinguished from neutral windows, with only 3% and 13% of hard or soft selective sweep miss-classified as neutral regions. Therefore, these performance analyses to discriminate among the different type of selective sweeps show that the method is sensitive and accurate under the present study design. The signals of selective sweeps identified in the LLP populations from the village and sylvatic area displayed a much higher number of windows classified as soft sweep and lower number of windows classified as hard sweep compared to the urban LBV population (Figure 4B) (*χ^2^* test=153.99, *p-value* < 1e^-5^). Such result highlights different modes of local adaptation acting on the urban population compared to those from the LLP National Park: a few *de novo* mutations was selected in the LBV city due to the strong selection pressure apply by insecticide, while positive selection on SGV is the most frequent mode of adaptation across less anthropized population in the La Lopé National Park.

### Candidate genes involved in local adaptation along the anthropogenic gradient

We obtained little overlap between the selection scans (both *XP-EHH* and *F_ST_*) and *DiploS/HIC* because the latter required large genomic windows of 110kb to apply the CNN classifier. In contrast, *XP-EHH* and *FST* can be conducted in relatively small genomic windows, or even at the site level. Nevertheless, we found that most of the selective sweeps detected by haplotype-based, frequency-based, and *DiploS/HIC* analyses in the LBV urban population were also identified as hard-(or linked-hard) selective sweeps. Overlap, the selection signals identified in the LBV urban population between the two types of analyses mainly focused on the known insecticide resistance and immune-related genes. In contrast, evidence of positive selection between in the village and sylvatic LLP populations were primarily soft-(or linked-soft) selective sweeps scattered along the autosomes (Figure 4). The 44 outlier regions identified by the haplotype-based *XP-EHH* statistic included a total of 118 candidate genes (Figure 4, Supplementary Table 4). Although gene ontology (GO) enrichment analysis did not show any significant over-or under-represented gene functions, some genes stood out from the group. The metal response element-binding Transcription Factor-1 (*MTF-1*, AGAP007377) was found to show clear signal of selection in LLP village, but not in the sylvatic LLP populations. This gene encodes for a zinc finger protein involved in the detoxification of non-essential, toxic heavy metals, such as cadmium, mercury, and silver. In parallel, in the sylvatic LLP population, the gene *pgant5* (AGAP012256) responsible for post-translational modification was found under strong positive selection in both comparisons with the village and the city. In *Drosophila*, this gene is a member of a large gene family known to participate in many aspects of development and organogenesis, notably in the upkeeping of a proper digestive system acidification (Tran et al. 2012).

## Discussion

Understanding the evolutionary mechanisms by which population and species adapt locally along heterogeneous habitats is fundamental to the evolution of diversity. Here, we evaluated how and by which population genetic processes a highly anthropophilic malaria-vector mosquito species, such as *An. coluzzii*, is able to colonize and adapt along an environmental gradient of anthropization in Gabon, including a forested area deprived of any permanent settlement in the La Lope National Park. To this end, we used a combination of population genetic approaches to analyze 77 whole-genome sequences of *An. coluzzii*, distributed along an anthropogenic gradient (urban, village and sylvatic areas). We investigate the demographic history of these Gabonese *An. coluzzii* populations in perspective with others from The Ag1000G Consortium (2020) across the species distribution range, as well as the selective forces driving local adaptation to the different habitats.

### An. coluzzii from Gabon exhibit singular genetic signatures

Consistent with previous studies (Pinto et al. 2013; Anopheles gambiae 1000 Genomes Consortium et al. 2020; Campos et al. 2021), we identified three major genetic groups across *An. coluzzii’*s distribution range: (i) the *West African* group composed of the mosquitoes from Burkina-Faso, Guinea, Ivory Coast, and Ghana; (ii) the Central African group composed of the Gabonese populations; and (iii) the South-Western African group with Angola (Figure 1C). These clusters coincide with the transitions between the central African rainforest belt and the Western northern and southern savannah biomes (Pinto, et al. 2013; Tene Fossog, et al. 2015). The Congo rainforest block has been suggested to constitute a potent barrier to gene flow (Pinto, et al. 2013; Tene Fossog, et al. 2015). Previous works from Campos, et al. (2021) showed that the Gabonese specimens clustered with those from the neighboring country, Cameroon, supporting a Central African cluster of *An. coluzzii* distinct from the others. Our analyses underlined the distinctiveness of the Gabonese LLP mosquitoes from more southern populations in Angola. Nevertheless, population-graph analysis suggested that coastal populations (i.e Libreville and Angola, Figure 1E) shared together a more recent common ancestor than with the West African group, north of the Congo-basin. This result is in agreement with previous works that evidence a subdivision between coastal and inland populations of *An. coluzzii* (Slotman et al. 2007; Tene Fossog et al. 2013). Moreover, the genetic characterization of the three populations from Gabon revealed higher level of homozygosity with regard to Western populations, consistent with smaller and more isolated populations (Figure 2). This result confirms that even at the “local” scale, the transition between coastal to rainforest area is associated with strong changes in the genetic make-up of *An. coluzzii* mosquitoes, as we can observe between Libreville and La Lope that are only ∼250km away from each other (Figure 1D)

### Historical rainforest evolution drives human and mosquitoes expansion in Central Africa

We estimated the split time between the Gabonese populations between the coastal urban area of LBV and LLP (rural/forested area) at *ca.* 8k ybp, followed by a secondary contact *ca.* 4.4kyr (Figure 3D). The initial split coincides with the peak of the African humid period (11 - 8kyr bp) and the major expansion of the African rainforest toward the tropics (Malhi et al. 2013). The Central African vegetation zones extended much further north (up to 400 – 500 km), and the Sahara was crisscrossed by lakes, rivers and inland deltas (Willis et al. 2013). These climate changes likely impacted human movements between Central African and other human African populations (Patin et al. 2009; Verdu et al. 2009; Batini et al. 2011; Lopez et al. 2018; Laval et al. 2019; Lopez et al. 2019), s well as the population dynamic of *Anopheles* mosquitoes. Interestingly, the secondary contact that restored gene flow between the coastal LBV population with the inland LLP population matches precisely the timing of the large-scale human expansion of the Bantu speaking people(Patin et al. 2009; Verdu et al. 2009; Batini et al. 2011; Lopez et al. 2018; Laval et al. 2019; Lopez et al. 2019). Fueled by agriculture, Bantu speaking human populations migrated most likely through the Central African tropical rainforest around 4,400 y ago, spreading the Bantu culture toward Central and South Africa (Koile et al. 2022). This period corresponds to the probable human specialization of *An. gambiae* and *An. coluzzii* (White et al. 2011; Ag1000G Consortium et al. 2020). It is thus very likely that large-scale human movements facilitated secondary contact between isolated mosquito populations (Figure 3D). The asymmetric gene flow since secondary contact, predominantly from the coastal into inland populations area may also reflect the constraints to thrive and develop in forested areas for *An. coluzzii* (Ayala et al. 2009; Tennessen et al. 2021). The post-glacial expansion (20k years ago) of the Bantu speaking farmers, and their colonization toward Central and South Africa (4000 to 5000 yr ago) likely reconnected isolated mosquito populations from the coastal and inland areas. In agreement with a scenario, La Lopé National Park has in fact a rich history of ancient human trades and migration corridor along the Congo Basin for Pygmy tribes, Bantu, and other people (UNESCO World Heritage Centre 2016), therefore, it could be fueled the movement and connection of human specialized mosquitoes. Also consistent with this scenario is the tempo of emergence within the genome of the forest-dueling Pigmy’s tribes of Central Africa of the genetic mutation that causes sickle-cell anaemia (also known as drepanocytosis) but also provides a strong selective advantage against malaria. This mutation emerged at least 20k year ago in Africa primarily in the genome of the Bantu’s farmers, and was only introduced ∼4000 to 5000 years ago into the rainforest pigmy hunter-gatherers of Central Africa, after admixture event between the two (Laval, et al. 2019).

### Urbanization exerts strong selection pressures in An. coluzzii

In contrast to the previously reported “shallow to moderate” population substructure reported among *An. coluzzii* populations within the West African group (*F_ST_* ≤ 0.04, Supplementary Figure S6) (The Ag1000G Consortium 2017, 2020), we observed moderate to high genetic structure within the Central African cluster at short geographic distance (*F_ST_* ≈ 0.05; Supplementary Figure S6). The urban coastal mosquito population from LBV was genetically highly distinct from LLP populations located only 250km inland. These urban mosquitoes displayed a population dynamic and demography that contrasted strongly with the population from the more natural environment of LPP (Figure 2). The urban LBV population was more inbred, it displayed stronger linkage disequilibrium, higher autozygosity, and a three-fold reduced effective population size compared to those from LLP. Such genetic contrasts likely reflect strong selection pressures to the highly anthropized urban environment, possibly induced by polluted larval habitats (Longo-Pendy et al. 2021). Consistent with this hypothesis, we found few but strong selection signals identified in the genome of the LBV populations, mostly (if not only) composed of hard-selective sweeps at sites well-known for insecticide resistance and detoxification processes (Figure 4A, Supplementary Table 4). Among them, we identified the knockdown resistance (*kdr*) mutations in the voltage-gated sodium channel (*VGSC*) of mosquitoes conferring resistance to pyrethroid insecticides (Martinez-Torres, et al. 1998; Ranson, et al. 2000; Davies, et al. 2007; Dong, et al. 2014; Clarkson, et al. 2021); the Rdl - GABA-gated chloride channel subunit gene, a locus with prior evidence of recent positive selection and/or an association with dieldrin insecticide resistance in *Anopheles* mosquitoes (Du, et al. 2005; Grau-Bové, et al. 2020); and the GSTE – Glutathione S-transferase epsilon gene cluster (Mitchell, et al. 2014). The LBV populations also harbored strong selective signal at the toll-like receptor signaling pathway known to mediate anti-pathogen defense, including against *Plasmodium* (Clayton et al. 2013). The African cities have been spared of *Anopheles* until recently (Robert et al. 2003). During the last two decades, *An.* coluzzii has overcome the constraints linked to breeding site pollution and it is now the predominant *Anopheles* species in the cities of Central Africa (Tene Fossog et al. 2015; Kamdem et al. 2017; Doumbe-Belisse et al. 2021; Longo-Pendy et al. 2021). As we observe in our study, this adaptation is associated to detoxication and insecticide resistant genes as previously documented (Antonio-Nkondjio et al. 2015; Kamdem et al. 2017). Therefore, urbanization tends to favor mosquito populations that are resistant to pollution and insecticides, which become less affected by actual vector control measures. Interestingly, the fact that a toll-like immune receptor is also strongly selected raises the question about how urban adaptation and pollution can affect vector competence to *Plasmodium*, a key aspect to understand and prevent urban malaria (Venkatesan 2024).

Given the asymmetric gene flow from LBV into LLP, we might have expected to find insecticide resistance alleles in the genomic background of the LLP population. However, no such selection signal was observed. Previous studies reported that mosquitoes carrying these insecticide resistance alleles display increased metabolism, and reduce body size, leading to reduced individual fitness of the mosquito that are not exposed to insecticide treatments (Oliver and Brooke 2016; Ingham, et al. 2017; Ingham, et al. 2021; Lucas, Nagi, Egyir-Yawson, Essandoh, Dadzie, Chabi, Djogbénou, et al. 2023; Romero, et al. 2023). For instance, such fitness effect led insecticide alleles to disappear in mosquitoes colonies (Ingham, et al. 2021). In La Lope village, no vector control strategies are implemented. Therefore, our results may indicate that resistant alleles are currently not detected, based on our limited sample size, but resistant alleles may be present at low frequency in the genomic background of LLP populations or they could swiftly appear as soon as insecticide selection pressure occurs by the gene flow with neighboring populations, including LBV.

### Standard genetic variation is key to local adaptation

In contrast to the coastal urban population of LBV, *An. coluzzii* inhabiting the LLP village and sylvatic areas displayed higher genetic diversity, slightly lower Tajima’s D, lower autozygosity and LD in their genomes. These genetic features suggest that the LLP populations have remained more stable in size for a longer time, and without any major bottleneck in a recent past, like those suspected in the LBV population in response to adaptation to polluted urban environment (Figure 2). These LLP populations are likely connected to other forested populations, across villages and unsampled sylvatic places. Despite the differences in hosts or breeding sites between both habitats at La Lope, the mosquitoes from the village and the sylvatic area were part of a same genetic pool (Figure 1C, Supplementary Figure S4). All analyses conducted to assess population genetic structure and differences in genetic diversity failed to discriminate mosquitoes from these two locations at putatively neutral and unlinked genetic markers (Supplementary Figure S6 and S7). Nevertheless, at this short geographic scale (∼10-15 Km) we expect a common genetic pool based on the potentially high dispersion rates in Anopheles (Costantini et al. 1999; Ayala et al. 2013; Smith et al. 2023). In fact, strong Isolation by distance (IBD) was previously detected in *An. coluzii* in other parts of Africa such as in the NW African Savanah, and major biogeographic barriers can also restrict dispersal of mosquitoes, such as the transition between Savanah and the rainforest biome of the Congo Basin (Lehmann et al. 2003; Pinto et al. 2013; Anopheles gambiae 1000 Genomes Consortium et al. 2020). Nevertheless, Battey et al (2020) showed that geographic location of individual *An. coluzzii* mosquito could be predicted from genetic data of the Ag1000G project with a precision of 5 km in average, with a median distance of 36 km (see also (Smith et al. 2023)). Thus, panmixia should be expected at such a small geographic scale like 10km, even if IBD may be strong in this species, and unless other processes are at play (e.g. selection).

The selection regime between domestic and sylvatic populations at La Lopé was dominated by soft-(or linked-soft) selective sweeps on the SGV (Figure 4), in contrast to a selective regime dominated by few hard-selective sweeps at LBV. The mode of adaptation of the LLP populations mostly involved selection on the SGV (or rapidly recurring beneficial mutations appearing on different haplotypes) (Harris, et al. 2018; Stephan 2019; Charlesworth and Jensen 2021; Johri, et al. 2022). This mode of adaptation has been shown to dominate also in other species like *Drosophila*, human and other organisms, even if some debates persist (Schrider and Kern 2017; Harris, et al. 2018; Stephan 2019; Charlesworth and Jensen 2021; Feder, et al. 2021; Garud, et al. 2021; Johri, et al. 2022). This predominance of soft (or link-soft) sweeps observed between LLP sylvatic and domestic populations was also shown to be relatively common across the genome of *An. coluzzii* (Xue et al. 2021). By analyzing the data from The Ag1000G Consortium (2017), Xue et al. (2021) reported that soft and partial selective sweeps were common place in the genome of 8 populations of *An. gambiae* and *An. coluzzii* across Africa. Nevertheless, mosquitoes from both LLP localities exhibited striking differences in selective signals marked by distinctive contrast at extended haplotype homozygosity (XP-EHH) scattered across their genomes. Noteworthy, no major differences in F_ST_ values were observed at these candidate genomic regions identified with elevated XP-EHH values (Figure 4A and Supplementary Figures 9 and 10). This discrepancy between XP-EHH and F_ST_ results further stress that selection signals are very likely soft or partial selective sweep involving only subtle changes in allele frequencies.

These selective signals scattered across the genomes did not reveal major genetic enrichment signals. As noted by Xue, et al. (2021) for the populations from The Ag1000G Consortium (2017), the numerous calls of selective sweep detected may reflect complex selective dynamics at play, for example, polygenic and quantitative trait adaptation (Pritchard, et al. 2010; Booker, et al. 2017), balancing selection (Connallon and Clark 2013), and introgression of beneficial alleles from neighboring unsampled populations (The Ag1000G Consortium 2017, 2020). Local adaptation of the mosquitoes to an un-anthropized habitat such as the sylvatic areas, compared to the village of the LLP National Park, likely involved many phenotypic traits that require further investigations. Nevertheless, such local adaptation are expected to be highly polygenic (Pritchard, et al. 2010) and is unlikely that all alleles involved are newly mutated.

Our study has evidenced the extraordinary ability of a major malaria vector to adapt to distinct ecological settings. In Central Africa, *An. coluzzii* inhabits from the most anthropogenic and polluted habitats to rural and wild protected natural areas such as National Parks. Indeed, this study characterized for the first time a population of a major malaria mosquito able to colonize and thrive in areas deprived of permanent human settlements, yet in direct vicinity of a village. Therefore, our results provide new clues about how mosquitoes adapted to humans while relying primarily on selection on their ancestral genetic polymorphism. Urban malaria is considered a major threat for malaria control and eradication in the coming years (Venkatesan 2024). However, protected areas have been relegated and little is known about how it can affect vector controls strategies, for instance, being a refuge for malaria vectors. Further insights are required on phenotypic changes, ecological and behavioral, of this and others malaria vectors, under this unique ecological scenario.

## Material and Methods

### Experimental design

Field mosquitoes were collected at the National Park of La Lopé, the village of La Lopé, and the city of Libreville between September 2015 and November 2017 (Figure 1A and Supplementary Table 1), according to the research permits required by the Gabonese government (AR0015/15/MESRS/CENAREST/CG/CST/CSAR) and the National Parks Agency (AE15011/PR/ANPN/SE/CS/AEPN). In the sylvatic and rural habitats of La Lopé National park, adult mosquitoes were sampled using Human Landing Catches (with the approval of the National Ethical Committee from Gabon PROT N° 0031/2014/SG/CNE) and BG traps (Biogents). Samples from Libreville city were collected by larvae dipping. A total of 96 mosquito samples were used in this study including 32 mosquitoes for each site (Supplementary Table 1).

### DNA extraction, genomic library preparation, whole genome sequencing

Total genomic DNA was extracted using DNeasy Blood & Tissue Kit (Qiagen) following the manufacturer’s instructions. DNA quality and concentration were estimated via PicoGreen (Promega). Genome library preparations took place at the Broad Institute using a Nextera XT Library Preparation Kit (Illumina). Libraries were sequenced on an Illumina HiSeq X instrument using a 300 cycle run format (150bp paired end reads).

### Bioinformatic data processing

To ensure compatibility of our data with those from the *Ag1000G* (Ag1000G Consortium 2017; Ag1000G Consortium et al. 2020), we followed the same bioinformatics protocols for SNP calling. Briefly, short reads were mapped to the *An. gambiae AgamP4* PEST reference assembly (Holt et al. 2002; Sharakhova et al. 2007) using *bwa-mem* version 0.7.17 (Li and Durbin 2009) with default parameters. Individuals with an average genome coverage depth lower than 14x (n=9) were excluded from downstream analyses (Supplementary Figure 1A and Figure 3A). After removing PCR duplicates with *Picard* tools (Anon 2019) and performing INDEL realignment with GATK’s *IndelRealigner* version 3.7 (McKenna et al. 2010), SNPs were called using GATK *Unified Genotyper* version 3.7. Low-quality SNP calls were filtered by removing variants that failed any of the following hard filters: QD<5, FS>60 and ReadPosRankSum < -8. We also retained only variants located within 63% of the genome previously classified as accessible in the Ag1000G consortium (Ag1000G Consortium 2017; Ag1000G Consortium et al. 2020).

Variants were then annotated using *snpEFF* version 4.3 (Cingolani 2022) with default parameters. Lastly, we generated a final high-quality SNP dataset using *vcftools* version 0.1.16 (Danecek et al. 2011) considering only biallelic SNPs, with genotypes with a genotype quality (GQ) higher than 20, discarding any variants with a missingness rate over 5% (*l-miss*<5%) and individuals (n=9) with a missingness rate over 10% (*i-miss*<10%).

The degree of relatedness among individuals (kinship coefficient) was estimated using *plink* version 1.9 (--genome) (Purcell et al. 2007). Haplotype estimation, also known as statistical phasing, was performed using *SHAPEIT2* version v2.r904 (Delaneau et al. 2008) with information from the reads, the reference haplotype panel of the Ag1000G phase-2 (ftp://ngs.sanger.ac.uk/production/ag1000g/phase2/AR1/haplotypes/), an effective population size (*N*e) of 1,000,000, default MCMC parameters and a window size of 2 Mb. Estimation of the ancestral versus derived allelic states of the SNPs was determined using an outgroup species. Following the Ag1000G consortium (Ag1000G Consortium et al. 2020), we polarized the SNP dataset using the consensus alleles defined from 10 *An. merus* from (Fontaine et al. 2015b). Polarized and phased datasets were composed respectively of a total of 5,859,776 and 2,982,164 SNPs for the whole genome (Supplementary Figure 3).

### Population genetic structure

To explore population structure in a larger, continent-wide context, we merged our Gabonese SNP dataset with the published phase-2 data from the Ag1000G project, considering only *An. coluzzii* species which include populations from Angola, Ivory Coast, Ghana, Guinea, and Burkina-Faso (The Anopheles gambiae 1000 Genomes Consortium 2020). Joint analyses between samples from Gabon and *An. coluzzii* samples from the Ag1000G phase-2 were performed by merging both VCF and keeping only SNPs at the intersection of both datasets. Following the Ag1000G Consortium (2017) methodologies, we investigated population genetic structure considering only the biallelic SNPs of the euchromatic freely recombining regions of chromosome 3, avoiding the peri-centromeric regions, and also avoiding well-known inversions on chromosome 2, heterochromatic regions, and the sexual X chromosome. From the regions 3R: 1-37 and 3L: 15-41 Mb, we removed SNPs in linkage disequilibrium, excluding SNPs above an *r^2^* threshold of 0.01 in moving windows of 500 SNPs with a step size of 250 SNPs via *scikit-allel* version 1.3.3 (Miles and Harding 2016). A minor allele frequency (MAF) of >1% was also applied. We first visualized population genetic structure using a PCA on the 1,003,463 unlinked SNPs using *scikit-allel*, considering genomic data from the Gabonese mosquito samples alone and combined with the data from the Ag1000G. Then, we quantified individual genetic ancestry proportions using the program *ADMIXTURE* version 1.3.0, testing various numbers of clusters (K) ranging from 2 to 10. *ADMIXTURE* was run for each K-value a 100-time, in the form of 10 replicate x 10 datasets, with each dataset composed of a random sampling of 100,000 variants from the total number of unlinked SNPs dataset. The most likely number of ancestral populations (K) was determined using the CV error rate (Alexander et al. 2009). Athough, the lowest CV error rate was obtained for *ADMIXTURE* models with K=3 ancestral populations, we found that further population sub-division was clearly recovered in simulations allowing up to K=6. These had clear support from other analyses (PCA, pairwise average *F_ST_*) and also from the previous study by the Ag1000G consortium (2020). From the 100 ADMIXTURE runs for each K, we use *CLUMPAK* version 1.1 with default settings to compare solutions and produce major and minor clustering solutions. In parallel, average *F_ST_* values were computed between all pairs of 8 populations, using the Hudson’s *F_ST_* estimator (Bhatia et al. 2013) with standard error for each average computed using a block-jackknife procedure in *scikit-allel*. *P-values* were estimated from the z-score following The Ag1000G consortium (2017; 2020).

To further explore the genetic relationship among populations, we performed an admixture graph analysis using *AdmixtureBayses* (Nielsen et al. 2023). Graphs were estimated using the pruned and polarized dataset of which individuals from Guinea were excluded due to their small sample size (n=5). We ran three independent MCMC chains each consisting of 22,500,000 steps (-n 450000), discarding the first 50% as burn-in. All other parameters were left as default. Finally, to test whether rural and sylvatic LLP samples formed a single panmictic population, we compared the observed versus the permuted jSFS using *δaδi* version 1.6.3. Permuted jSFS was created using *δaδi* build-in function “scramble_pop_ids” from the dadi.Spectrum_mod which generates an average spectrum expected overall permutation of the individuals in the dataset.

### Genetic diversity

We quantified the level of genetic diversity in each population by computing several descriptive statistics from chromosome arms 3L and 3R excluding the pericentromeric regions with *scikit-allel*. Nucleotide diversity (π) and Tajima’s D were calculated in 10-kb non-overlapping windows. Runs of homozygosity (ROH) were defined as contiguous regions of an individual’s genome where only homozygous blocks were identified through a HMM function implemented in *scikit-allel* as described in the (Anopheles gambiae 1000 Genomes Consortium *et al*, 2017). Following the AG1000G, all the autosomes were used for this analysis. Tracts of identity-by-descent (IBD) between all pairs of individuals within each of the 8 populations were inferred by IBDseq version r1206 (Browning and Browning 2013) with default parameters. Folded SFS (a.k.a. the minor allele frequency spectrum) was computed using allele counts using *scikit-allel*. To facilitate comparison with theoretical SFS for a population with constant size (expected to have the constant scaled frequency for all values of k), we scaled each folded SFS by a factor (k * (n – k) / n) where k is the minor allele count and n is the number of chromosomes following the Ag1000G (2017). LD decay was computed by calculating the genotype correlation coefficient *r^2^* (Rogers & Huff, 2009) for randomly sampled pairs of SNPs at distances raging from 10 to 10^7^ bp using *scikit-allel*.

### Demographic history

We estimated the long-term demographic history of each population from Gabon using *Stairway plot* v.2 (Liu and Fu 2020). This approach estimates *Ne* variation back to the time of the most recent common ancestor (TMRCA) based on the unfolded SFS. The full unfolded SFS was generated for each population using *scikit-allel* from the peri-centromeric euchromatic regions of chromosome 3 of the polarized SNP dataset (Anopheles gambiae 1000 Genomes Consortium *et al*, 2017). To translate *Stairway plot* estimates of *Ne* and time into natural units (i.e., individuals and years respectively) we assumed a generation time of 11 per year and a mutation rate of 3.5 x 10^-9^ per bp (Keightley et al. 2014) and per generation following the Ag1000G (Ag1000G Consortium 2017).

We used *δaδi* (Gutenkunst et al. 2009) to infer the best fitting demographic model of population isolation, possibly with migration, between populations from LBV and LLP village using the polarized SNPs dataset on chromosome 3, without any LD-pruning nor any MAF filtering. We considered a total of eight alternative nested models of historical divergence, which were built on four basic population isolation models: (i) a model of Strict Isolation without gene flow (SI), (ii) a model of Isolation with continuous Migration (IM), (iii) a model of divergence with initial migration or Ancient Migration (AM), and (iv) a model of Secondary Contact (SC). These models were extended to integrate temporal variation in effective population size (G), enabling exponential growth or contraction.

For each population, the SFS was computed for a number of individuals projected onto the smallest population sample size (N=20 diploid samples for LLP village). A joined SFS (jSFS) between the pair of populations was generated for the sites in the genome that did not contain any missing data. The 8 models were fitted independently using successively a hot and a cold simulated annealing procedure followed by “BFGS” optimization (Tine et al. 2014). We set the grid points to {n, n + 10, n + 20}, where “n” is the number of haploid chromosomes (n=40). Model parameter bounds for *Ne* scalars were *N* **∈** (0.01, 100), for the population exponential growth parameter *b* **∈** (0.01, 100), for the time were *T* **∈** (0, 10), for migration were *m* **∈** (0, 50), and for the genotyping uncertainty *O* **∈** (0.01, 0.99). We ran 100 independent optimizations for each model to check for convergence and retrained the best one. Comparisons among models were based on the Akaike information criterion (AIC). We use the framework developed by (Rougeux et al. 2017) to address over-parametrization issue and to penalize models containing more parameters. We used a conservative threshold to retain models with ΔAIC<10. This procedure identified the best-fitted model to our data and involved a population isolation with secondary contact and exponential population size change (SCG). Model parameters were converted into natural units as follows: ancestral effective population size (Ne) was calculated by *Ne* = *θ*/(4.*µ*.*l*), where *θ* is the scaled population mutation rate (*θ*=4.*Ne*.*µ*.*l*), *µ* is the mutation rate per site and per generation (*µ* = 3.5 x 10^-9^), and *l* the length of the analyzed sequence (*l* = 39,359,290). The effective population size of populations 1 and 2 are given in units of *Ne1 = nu1 x Nref* and *Ne2 = nu2 x Nref*, where *nu1* and *nu2* are the population size relative to the size *Nref* of the ancestral population. Estimation of times in units of 2*Nref* generations (*Ts* and *Tsc*) were converted into years assuming a generation time of 0.09 years (equal to 11 generation per year). Estimated migration rates (*m12* and *m21*) were divided by 2*Nref* to obtain the proportion of migrants received by each population every generation. The number of migrants per generation were obtained by *Nref* x *nu1* x *m12* and *Nref* x *nu2* x *m21*. To estimate parameter uncertainty, we used the Godambe information matrix method from *δaδi*. Nonparametric bootstrapping was used to generate 1,000 bootstrapped data sets to estimate the 95% confidence intervals (CIs) using the standard error of maximum likelihood estimates (se) (Supplementary Table 3).

### Identification of selection signatures

To identify candidate genes and genomic regions impacted by selection histories that varied geographically between sylvatic, rural, and urban areas, we first compared allele frequencies and haplotype diversity between the sampling sites. Genome scan plots of between-population statistics were computed using *scikit-allel* to report the Hudson’s *F_ST_* estimator statistic and the average nucleotide difference between pairs of populations (Dxy). These statistics were computed in blocks of 1000 SNPs along the genome. To detect long stretches of homozygosity in a given population relative to another population (Sabeti *et al*, 2007) we estimated the XP-EHH using the *R* package *rehh* version 3.2.2 (Gautier et al. 2017) using the maxgap=20kb option to limit the extension of a haplotype through a gap of 20kb. *P*-value associated to each SNP was adjusted for the false discovery rate (FDR) with a threshold of 0.05. Candidate regions were identified using the R function *calc_candidate_regions* from the *rehh* package with a minimum number of significant SNPs in the region equal to 3 (min_n_extr_mrk=3) and a windows size of 10kb (window_size=1e4). All genes spanning the candidate regions were reported in the Supplementary Table 4.

### Deep learning classification of genomic windows to identify categories of selective sweeps

We used a deep learning approach based on the convoluted neural network (CNN) implemented in *diploS/HIC* (Kern and Schrider 2018) to classify genomics windows into five categories of selective sweep: hard sweep (or linked hard), soft sweep (or linked soft), or neutral (as a null hypothesis). Using the coalescent based simulator of *discoal* (Kern and Schrider 2016), we first generated simulations of 110kb genomic regions under the different categories of selection tested. These genomic regions were then further split into 11 sub-regions of 10kb to allow the CNN classifier capture the genomic properties of the windows neighboring the central focal window. The training and test data sets were produced using the same properties and demographic histories as observed in each of our three Gabonese populations under the five different selection scenarios. The demographic history generated by *Stairway plot2* for each population was used to generate simulated datasets as realistic as possible to our populations. A total of 2,000 training genomic regions and 1,000 testing regions with a single sweep were generated for each population, using as simulation parameters a per-site and per generation mutation rate of 3.5e-9 and 11 generations per year.

Once simulated, we first investigated the goodness-of-fit of the simulations to our observed data, and the prediction performance and accuracy of each of the model considered. These are critical steps to determine whether the model is well-trained and to assess decisions that could be made from poorly fitting models (A. Kern, *pers. comm.)*. We assessed the goodness-of-fit between the simulated and empirical data by visualizing the distribution of the observed values for each of the 12 descriptive statistics used by *DiploS/HIC* compared to the simulated distributions obtained for each type of selection (Supplementary Figure S11). For each descriptive statistic, a good convergence was observed between simulated and empirical data, with the only noticeable exception observed for the nucleotide diversity (π), which displays a reduced amount of diversity in our simulated dataset regardless of the population of interest. Then, the performance and accuracy of the CNN classifier for each model of selection were estimated using the confusion matrix (Supplementary Figure S12). In addition, samples from the LPV were found to exhibit a slight but significantly higher accuracy than the population from LBV and the LLP sylvatic area, conserved across all sweep types, indicating a better classification performance. In order to assess the convergence of the results, we generated 10 different coalescent simulated datasets for each population and each dataset was used to train 10 times the CNN classifier, resulting in a total of 100 runs. In order to compare and interpret the convergence between the predictions of each run, we first filtered out selective sweep having a low probability of being neutral (p>0.01) and considered only sweep observed in at least 50% of our 100 replicated runs.

## Acknowledgements

We thank the Ecology of Vectorial Systems team at the CIRMF (Franceville, Gabon) for their support in field collections. We thank the volunteers from the La Lopé village (Gabon) for their collaboration and their compliance during the field investigations. We are especially grateful to the National Park Agency in Gabon (ANPN) for granting us access to the research station SEGC at the National Park of La Lopé. We thank the members of the Fontaine’s group and especially Margaux Lefebvre for sharing input and scripts for the *AdmixtureGraph* analysis. We also thank the Center for Information Technology of the University of Groningen for their support and for providing access to the HPC cluster. We also acknowledge the ISO-9001 certified IRD i-Trop HPC (https://bioinfo.ird.fr), member of the South Green Platform (www.southgreen.fr) at IRD Montpellier for providing HPC resources that have contributed to the results reported in this paper. This study was supported by a grant from the ANR, France (ANR-18-CE35-0002-01 – WILDING) awarded to D.A. L.B was supported by the ARTS doctoral fellowship program from the IRD. DEN and JT were supported in part with federal funds from the National Institute of Allergy and Infectious Diseases, National Institutes of Health, Department of Health and Human Services, under Grant Number U19AI110818 to the Broad Institute. Finally, we thank Thee H. Bert for the valuable discussions.

## Author contributions

Conception: D.A. and D.E.; funding acquisition: D.A. and D.E.; biological data acquisition and management: D.A., L.B., N.R., B.M, O.A.E, M.F.N., N.M.L.P., C.P.; sequence data acquisition: D.E. and J.T.; method development and data analysis: J.D. and M.C.F.; interpretation of the results: J.D., M.C.F., and D.A.; drafting of the manuscript: J.D., M.C.F., and D.A.; reviewing and editing of the manuscript: J.D., M.C.F., and D.A., with inputs from all the co-authors.

## Data Availability

Raw sequencing data generated as part of this project are available under the Bioproject PRJNAXXXX. Biosample and SRA accession numbers are provided in Supplementary Table 1. Data, codes and related documentations that support the findings of this study are openly available in the DataSuds repository (IRD, France) at DOI: To Be Announced. Data reuse is granted under a CC-BY license. Source data are provided with this paper. Codes and related documentations that support the findings of this study are also openly available via GITHUB LINKS (https://github.com/jdaron/wilding). Data and code reuse is granted under a CC-BY license.

## Supplement

**Supplementary Figure 1:** (A) Workflow of the reads mapping and SNP genotyping procedures used. (B) Schematic overview of the different analysis and the input dataset used.

**Supplementary Figure 2: Sequencing coverage depth and SNP density along the genome.**

**(A)** Distribution of the mean sequencing depth of the 96 samples. Bars represent individual mosquito samples and are color-coded according to their sampling origin: Libreville (yellow); La Lopé village (blue); and La Lopé sylvatic (green). The horizontal line represents the coverage cut-off at 14x used to exclude samples below that threshold. **(B)** Coverage ratio of the mean sequencing depth for each chromosome over the coverage of the whole genome. This allowed to assign the sex of each individual, which was unknown for the larvae from Libreville (see Supplementary Table S1). **(C)** Density of the high-quality SNPs in 200-kb non-overlapping windows over the genome.

**Supplementary Figure 3: Kinship analysis across the Gabonese dataset estimated with pair-wise IBD estimator (PI_HAT) between samples in PLINK.** The threshold 0.1875 represents the half-way point between 2nd and 3rd degree relatives and is a common cut-off to use.

**Supplementary Figure 4: (A)** Scree-plot showing the variance fraction explained by each principal component of the PCA for the African *An. coluzzii* samples (combining the Gabonese and AG1000G datasets) represented in Figure 1. **(B)** PCA of the 77 *An. coluzzii* mosquitoes from Gabon retained for further analysis using biallelic SNPs from the euchromatic regions of the chromosome 3. The bar chart shows the percentage of variance explained by each principal component.

**Supplementary Figure 5: Analysis of population structure and genetic ancestry in *An. coluzzii* considering the Gabonese populations in perspective with those from the *Ag1000G*.** (A) Individual ancestry proportions (from K=2 to K=10) were estimated using the ADMIXTURE program. Each vertical bar represents an individual mosquito grouped according to sampling location and colored according to the proportion of the genome inherited from each of the *K* ancestral clusters tested. (B) Box-plots showing the average (red dots), median, and interquartile values of the cross-validation (CV) error rate estimated using the ADMIXTURE program for each ancestral cluster tested (with K ranging between 2 and 10). Black dots show the CV error rate values for 100 replicated runs at each tested K values. K=3 was chosen as the best-fitted solution for our SNP dataset, since that value minimizes the CV error rate.

**Supplementary Figure 6: Pairwise population differentiations (*F_ST_*) among populations of *An. coluzzii*.** Average differentiation in allele frequency estimated using the *F_ST_* statistics between pairs of populations. Upper left portion of the matrix shows average *F_ST_* values between each population pair. Bottom right portion of the matrix shows the *p-value* derived from the z-score for each *F_ST_* value estimated via a block-jackknife procedure.

**Supplementary Figure 7: Test of departure from random mating expectation (panmixia) between pairs of populations performed using *δaδi***. The left panels represent the observed joint site frequency spectrum (jSFS) between populations pairs along with a model fit and residuals using *δaδi*, for a “scramble” model where individuals are permuted across population. On the right panel, the null distribution of *χ*^2^ values is obtained by measuring the deviation between 1000 replicates of permuted individual labels across the population pair to the scramble model. Vertical black line and value correspond to the *χ*^2^ value calculated between the observed jSFS and the scramble jSFSmodel. This test of departure from panmictic expectation was performed for all population pairs including: La Lopé village versus La Lopé sylvatic (A), Libreville versus La Lopé village (B), and Libreville versus La Lopé sylvatic (C).

**Supplementary Figure 8: *δaδi* model selection based on the AIC score obtained for 8 different models with 100 replicates**. The lowest AIC score was observed for the model secondary contact with growth (SCG).

**Supplementary Figure 9: Genome scan of *F_ST_* and *D_xy_* statistics**. Both statistics were calculated in 1000kb non-overlapping windows and plotted for all pairwise comparisons. Fine horizontal black lines indicate the heterochromatic regions excluded from the analyses, and dotted lines indicate known insecticide resistance genes.

**Supplementary Figure 10: Genome scan of XP-EHH scores calculated at the SNP level along the genome and plotted for all pairwise population comparisons**. For each population comparison (e.g., LPV vs LBV), positive scores indicate longer haplotype homozygosity and therefore recent selection in the first population (e.g., LLP village), and negative scores indicate selection in the second population (e.g., LBV). Each dot has been colored by its associated *p-*value for the XP-EHH score, with shaded red and blue colors gradient representing non-significant SNPs, and red and blue representing significant SNPs (p-value < 1e^-4^). Gray areas above the x-axis indicate heterochromatic regions excluded from our analysis, and dotted lines indicate candidate genes including known insecticide genes.

**Supplementary Figure 11**: (A) Barplot representing the proportion of SNPs displaying significant *p*-value for the XP-EHH score for each of the 3 pairwise population comparisons. Color code represents the population in which the SNP has been found significant (Yellow – LBV; Green – LLP sylvatic; Blue – LLP village). (B) Functional annotation of the SNPs identified as significant using the XP-EHH scores in each of the three pairwise comparisons.

**Supplementary Figure 12: Goodness-of-fit between empirical and simulated data under the 5 different types of selection scenarios of selective sweep in the *diploS/HIC* analysis**. The distributions obtained for each of the 12 summary statistics used in *diploS/HIC* and for each population (LBV, LPV, LPS) are displayed for the simulated (red) and empirical (blue) data.

**Supplementary Figure 13**: **Graphical representation of the confusion matrix.** Each matrix is represented in form of a barplot, with each facet of the figure represent the true label of each testing set, the x-axis represents the predicted type of selective sweep and the y-axis the proportion of windows assigned to each sweep type. Each barplot consist of the average of the 100 replicates with the confidence interval represented as error bar.

**Supplementary Table 1: Description of the *An. coluzzii* sampling from Gabon for which whole genome sequencing was performed and sequencing statistics.**

**Supplementary Table 2: Comparison among the 8 models of population isolation estimated with δaδi.** For each models the fittest model has been determined based on the lowest maximum likelihood (MLE) value. Comparisons among models were based on the value of Akaike information criterion (AIC) and the ΔAIC value. The optimized demographic parameter values for each of the 8 models tested were converted with the estimate of *Theta* (*θ*): the ancestral effective population size before population split (*Nref*); the effective population size after split for La Lopé rural (*nu1*) and Libreville (*nu2*) populations; the exponential growth coefficient for La Lopé rural (*b1*) and Libreville (*b2*) populations. The *b* parameter is defined as a ratio of contemporary to ancestral effective population size (ancestral meaning after splitting time). Population showing an exponential growth is associated with *bi*>1 and reduction in population effective size with *bi*<1. Migration parameters include migration rates from Libreville population into La Lopé rural population (*m_12_*) and reciprocally (*m_21_*). Time parameters include the duration (in years) of the allopatric divergence period (*T_split_*), and the duration of the migration period (i.e., *T_AM_* for the AM models and *T_SC_* for the secondary contact models). The parameter (O) is the proportion of correct SNP orientation. The estimated value of each parameter was converted so that migration rates represent the fraction of a population replaced by migrants every generation, and temporal parameters appear in years.

**Supplementary Table 3:** Details of the demographic parameter values estimated by δaδi for the best fit model of secondary contact with population size change (SC+G). The optimized demographic parameter values were converted with the estimate of *Theta* (*θ*): the ancestral effective population size before population split (*Nref*); the effective population size after split for La Lopé rural (*nu1*) and Libreville (*nu2*) populations; the exponential growth coefficient for La Lopé rural (*b1*) and Libreville (*b2*) populations. The *b* parameter is defined as a ratio of contemporary to ancestral effective population size (ancestral meaning after splitting time). Population showing an exponential growth is associated with *bi*>1 and reduction in population effective size with *bi*<1. Migration parameters include migration rates from Libreville population into La Lopé rural population (*m_12_*) and reciprocally (*m_21_*). Time parameters include the duration (in years) of the allopatric divergence period (*T_split_*), and the duration of the migration period (i.e., *T_AM_* for the AM models and *T_SC_* for the secondary contact models). The parameter (O) is the proportion of correct SNP orientation. The estimated value of each parameter was converted so that migration rates represent the fraction of a population replaced by migrants every generation, and temporal parameters appear in years. Parameters uncertainties and 95% confidence intervals were estimated based on the Godambe Information Matrix, computed from 100 bootstrapped data sets (uncert_GIM function of dadi. Godambe module).

|  | parameters | biological units | ci_low | ci_upp |
| --- | --- | --- | --- | --- |
| **nu1** | 6.08 | 865 159.02 | 0 | 6 054 973.88 |
| **nu2** | 2.05 | 292 460.51 | 0 | 7 667 286.19 |
| **b1** | 0.10 | 0.10 | 0.09 | 0.11 |
| **b2** | 0.31 | 0.31 | 0.25 | 0.37 |
| **m12** | 8.26 | 0.000029 | 0 | 8.26 |
| **m21** | 4.68 | 0.000016 | 0 | 4.68 |
| **Ts** | 0.30 | 7624.43 | 0 | 6 586 498.64 |
| **Tsc** | 0.17 | 4431.69 | 0 | 741 840.30 |
| **O** | 0.99 | 0.99 | 0.98 | 1.00 |
| **Nref** | 78 429.51 | 142 332.55 | 0 | 82 941 978.22 |

**Supplementary Table 4: Candidate regions and genes identified by the XP-EHH test from *rehh* program, and functional annotation of the significant SNPs included in those regions.**

## Notes

### Competing Interest Statement

The authors have declared no competing interest.

### Summary of Updates

The manuscript previously submitted to bioRxiv has been resubmitted following minor corrections, including typographical errors and similar adjustments.

## References

Ag1000G Consortium. 2017. Genetic diversity of the African malaria vector Anopheles gambiae. Nature 552:96–100.

Ag1000G Consortium CS, Miles A, Harding NJ, Lucas ER, Battey CJ, Amaya-Romero JE, Kern AD, Fontaine MC, Donnelly MJ, Lawniczak MKN, et al. 2020. Genome variation and population structure among 1142 mosquitoes of the African malaria vector species Anopheles gambiae and Anopheles coluzzii. Genome Res. 30:1533–1546.

Alexander DH, Novembre J, Lange K. 2009. Fast model-based estimation of ancestry in unrelated individuals. Genome Res. 19:1655–1664.

Anon. 2019. Picard toolkit. Broad Institute Available from: http://broadinstitute.github.io/pi-card/

Anopheles gambiae 1000 Genomes Consortium CS, Miles A, Harding NJ, Lucas ER, Battey CJ, Amaya-Romero JE, Kern AD, Fontaine MC, Donnelly MJ, Lawniczak MKN, et al. 2020. Genome variation and population structure among 1142 mosquitoes of the African malaria vector species Anopheles gambiae and Anopheles coluzzii. Genome Res. 30:1533–1546.

Antonio-Nkondjio C, Tene Fossog B, Kopya E, Poumachu Y, Menze Djantio B, Ndo C, Tchuinkam T, Awono-Ambene P, Wondji CS. 2015. Rapid evolution of pyrethroid resistance prevalence in Anopheles gambiae populations from the cities of Douala and Yaoundé (Cameroon). Malar J 14:155.

Ayala D, Acevedo P, Pombi M, Dia I, Boccolini D, Costantini C, Simard F, Fontenille D. 2017. Chromosome inversions and ecological plasticity in the main African malaria mosquitoes. Evolution 71:686–701.

Ayala D, Costantini C, Ose K, Kamdem GC, Antonio-Nkondjio C, Agbor J-P, Awono-Ambene P, Fontenille D, Simard F. 2009. Habitat suitability and ecological niche profile of major malaria vectors in Cameroon. Malaria Journal 8:307.

Ayala D, Guerrero RF, Kirkpatrick M. 2013. REPRODUCTIVE ISOLATION AND LOCAL ADAPTATION QUANTIFIED FOR A CHROMOSOME INVERSION IN A MALARIA MOSQUITO. Evolution 67:946–958.

Ayala D, Ullastres A, González J. 2014. Adaptation through chromosomal inversions in Anopheles. Front. Genet. [Internet] 5. Available from: https://www.frontiersin.org/journals/genetics/articles/10.3389/fgene.2014.00129/full

Ayala FJ, Coluzzi M. 2005. Chromosome speciation: Humans, Drosophila, and mosquitoes. Proceedings of the National Academy of Sciences 102:6535–6542.

Barrón MG, Paupy C, Rahola N, Akone-Ella O, Ngangue MF, Wilson-Bahun TA, Pombi M, Kengne P, Costantini C, Simard F, et al. 2019. A new species in the major malaria vector complex sheds light on reticulated species evolution. Sci Rep 9:14753.

Batini C, Lopes J, Behar DM, Calafell F, Jorde LB, van der Veen L, Quintana-Murci L, Spedini G, Destro-Bisol G, Comas D. 2011. Insights into the Demographic History of African Pygmies from Complete Mitochondrial Genomes. Molecular Biology and Evolution 28:1099– 1110.

Bhatia G, Patterson N, Sankararaman S, Price AL. 2013. Estimating and interpreting FST: The impact of rare variants. Genome Res. 23:1514–1521.

Browning BL, Browning SR. 2013. Detecting Identity by Descent and Estimating Genotype Error Rates in Sequence Data. The American Journal of Human Genetics 93:840–851.

Campos M, Hanemaaijer M, Gripkey H, Collier TC, Lee Y, Cornel AJ, Pinto J, Ayala D, Rompão H, Lanzaro GC. 2021. The origin of island populations of the African malaria mosquito, Anopheles coluzzii. Commun Biol 4:1–9.

Cingolani P. 2022. Variant Annotation and Functional Prediction: SnpEff. In: Ng C, Piscuoglio S, editors. Variant Calling: Methods and Protocols. Methods in Molecular Biology. New York, NY: Springer US. p. 289–314. Available from: 10.1007/978-1-0716-2293-3_19

Clayton AM, Dong Y, Dimopoulos G. 2013. The Anopheles Innate Immune System in the Defense against Malaria Infection. Journal of Innate Immunity 6:169–181.

Costantini C, Sagnon N, della Torre A, Coluzzi M. 1999. Mosquito behavioural aspects of vector-human interactions in the Anopheles gambiae complex. Parassitologia 41:209–217.

Danecek P, Auton A, Abecasis G, Albers CA, Banks E, DePristo MA, Handsaker RE, Lunter G, Marth GT, Sherry ST, et al. 2011. The variant call format and VCFtools. Bioinformatics 27:2156–2158.

Davies TGE, Field LM, Usherwood PNR, Williamson MS. 2007. A comparative study of voltage-gated sodium channels in the Insecta: implications for pyrethroid resistance in Anopheline and other Neopteran species. Insect Molecular Biology 16:361–375.

Delaneau O, Coulonges C, Zagury J-F. 2008. Shape-IT: new rapid and accurate algorithm for haplotype inference. BMC Bioinformatics 9:540.

Doumbe-Belisse P, Kopya E, Ngadjeu CS, Sonhafouo-Chiana N, Talipouo A, Djamouko-Djonkam L, Awono-Ambene HP, Wondji CS, Njiokou F, Antonio-Nkondjio C. 2021. Urban malaria in sub-Saharan Africa: dynamic of the vectorial system and the entomological inoculation rate. Malar J 20:364.

Fontaine MC, Pease JB, Steele A, Waterhouse RM, Neafsey DE, Sharakhov IV, Jiang X, Hall AB, Catteruccia F, Kakani E, et al. 2015a. Extensive introgression in a malaria vector species complex revealed by phylogenomics. Science 347:1258524.

Fontaine MC, Pease JB, Steele A, Waterhouse RM, Neafsey DE, Sharakhov IV, Jiang X, Hall AB, Catteruccia F, Kakani E, et al. 2015b. Extensive introgression in a malaria vector species complex revealed by phylogenomics. Science 347:1258524.

Garud NR, Messer PW, Buzbas EO, Petrov DA. 2015. Recent Selective Sweeps in North American Drosophila melanogaster Show Signatures of Soft Sweeps. PLOS Genetics 11:e1005004.

Gautier M, Klassmann A, Vitalis R. 2017. rehh 2.0: a reimplementation of the R package rehh to detect positive selection from haplotype structure. Molecular Ecology Resources 17:78–90.

Gutenkunst RN, Hernandez RD, Williamson SH, Bustamante CD. 2009. Inferring the Joint Demographic History of Multiple Populations from Multidimensional SNP Frequency Data. PLOS Genetics 5:e1000695.

Hermisson J, Pennings PS. 2005. Soft Sweeps: Molecular Population Genetics of Adaptation From Standing Genetic Variation. Genetics 169:2335–2352.

Holt RA, Subramanian GM, Halpern A, Sutton GG, Charlab R, Nusskern DR, Wincker P, Clark AG, Ribeiro JoséMC, Wides R, et al. 2002. The Genome Sequence of the Malaria Mosquito Anopheles gambiae. Science 298:129–149.

Kamdem C, Fouet C, Gamez S, White BJ. 2017. Pollutants and Insecticides Drive Local Adaptation in African Malaria Mosquitoes. Molecular Biology and Evolution 34:1261–1275.

Kawecki TJ, Ebert D. 2004. Conceptual issues in local adaptation. Ecology Letters 7:1225– 1241.

Keightley PD, Ness RW, Halligan DL, Haddrill PR. 2014. Estimation of the Spontaneous Mutation Rate per Nucleotide Site in a Drosophila melanogaster Full-Sib Family. Genetics 196:313–320.

Kern AD, Schrider DR. 2016. Discoal: flexible coalescent simulations with selection. Bioinformatics 32:3839–3841.

Kern AD, Schrider DR. 2018. diploS/HIC: An Updated Approach to Classifying Selective Sweeps. G3 Genes|Genomes|Genetics 8:1959–1970.

Klassmann A, Gautier M. 2022. Detecting selection using extended haplotype homozygosity (EHH)-based statistics in unphased or unpolarized data. PLOS ONE 17:e0262024.

Koile E, Greenhill SJ, Blasi DE, Bouckaert R, Gray RD. 2022. Phylogeographic analysis of the Bantu language expansion supports a rainforest route. Proceedings of the National Academy of Sciences 119:e2112853119.

Kyalo D, Amratia P, Mundia CW, Mbogo CM, Coetzee M, Snow RW. 2017. A geo-coded inventory of anophelines in the Afrotropical Region south of the Sahara: 1898-2016. Wellcome Open Res 2:57.

Laval G, Peyrégne S, Zidane N, Harmant C, Renaud F, Patin E, Prugnolle F, Quintana-Murci L. 2019. Recent Adaptive Acquisition by African Rainforest Hunter-Gatherers of the Late Pleistocene Sickle-Cell Mutation Suggests Past Differences in Malaria Exposure. Am J Hum Genet 104:553–561.

Lehmann T, Licht M, Elissa N, Maega BTA, Chimumbwa JM, Watsenga FT, Wondji CS, Simard F, Hawley WA. 2003. Population Structure of Anopheles gambiae in Africa. J Hered 94:133–147.

Li H, Durbin R. 2009. Fast and accurate short read alignment with Burrows-Wheeler transform. Bioinformatics 25:1754–1760.

Liu X, Fu Y-X. 2020. Stairway Plot 2: demographic history inference with folded SNP frequency spectra. Genome Biology 21:280.

Longo-Pendy NM, Tene-Fossog B, Tawedi RE, Akone-Ella O, Toty C, Rahola N, Braun J-J, Berthet N, Kengne P, Costantini C, et al. 2021. Ecological plasticity to ions concentration determines genetic response and dominance of Anopheles coluzzii larvae in urban coastal habitats of Central Africa. Sci Rep 11:15781.

Lopez M, Choin J, Sikora M, Siddle K, Harmant C, Costa HA, Silvert M, Mouguiama-Daouda P, Hombert J-M, Froment A, et al. 2019. Genomic Evidence for Local Adaptation of Hunter-Gatherers to the African Rainforest. Curr Biol 29:2926–2935.e4.

Lopez M, Kousathanas A, Quach H, Harmant C, Mouguiama-Daouda P, Hombert J-M, Froment A, Perry GH, Barreiro LB, Verdu P, et al. 2018. The demographic history and mutational load of African hunter-gatherers and farmers. Nat Ecol Evol 2:721–730.

Louis M, Galimberti M, Archer F, Berrow S, Brownlow A, Fallon R, Nykänen M, O’Brien J, Roberston KM, Rosel PE, et al. 2021. Selection on ancestral genetic variation fuels repeated ecotype formation in bottlenose dolphins. Science Advances 7:eabg1245.

Lynd A, Oruni A, van’t Hof AE, Morgan JC, Naego LB, Pipini D, O’Kines KA, Bobanga TL, Donnelly MJ, Weetman D. 2018. Insecticide resistance in Anopheles gambiae from the northern Democratic Republic of Congo, with extreme knockdown resistance (kdr) mutation frequencies revealed by a new diagnostic assay. Malaria Journal 17:412.

Ma Y, Ding X, Qanbari S, Weigend S, Zhang Q, Simianer H. 2015. Properties of different selection signature statistics and a new strategy for combining them. Heredity 115:426–436.

Malhi Y, Adu-Bredu S, Asare RA, Lewis SL, Mayaux P. 2013. African rainforests: past, present and future. Philosophical Transactions of the Royal Society B: Biological Sciences 368:20120312.

McKenna A, Hanna M, Banks E, Sivachenko A, Cibulskis K, Kernytsky A, Garimella K, Altshuler D, Gabriel S, Daly M, et al. 2010. The Genome Analysis Toolkit: a MapReduce framework for analyzing next-generation DNA sequencing data. Genome Res. 20:1297–1303.

Messer PW, Petrov DA. 2013. Population genomics of rapid adaptation by soft selective sweeps. Trends in Ecology & Evolution 28:659–669.

Miles A, Harding N. 2016. scikit-allel: A Python package for exploring and analysing genetic variation data.

Munhenga G, Brooke BD, Spillings B, Essop L, Hunt RH, Midzi S, Govender D, Braack L, Koekemoer LL. 2014. Field study site selection, species abundance and monthly distribution of anopheline mosquitoes in the northern Kruger National Park, South Africa. Malar J 13:27.

Nielsen SV, Vaughn AH, Leppälä K, Landis MJ, Mailund T, Nielsen R. 2023. Bayesian inference of admixture graphs on Native American and Arctic populations. PLOS Genetics 19:e1010410.

Patin E, Laval G, Barreiro LB, Salas A, Semino O, Santachiara-Benerecetti S, Kidd KK, Kidd JR, Veen LV der, Hombert J-M, et al. 2009. Inferring the Demographic History of African Farmers and Pygmy Hunter–Gatherers Using a Multilocus Resequencing Data Set. PLOS Genetics 5:e1000448.

Paupy C, Makanga B, Ollomo B, Rahola N, Durand P, Magnus J, Willaume E, Renaud F, Fontenille D, Prugnolle F. 2013. Anopheles moucheti and Anopheles vinckei Are Candidate Vectors of Ape Plasmodium Parasites, Including Plasmodium praefalciparum in Gabon. PLOS ONE 8:e57294.

Pinto J, Egyir-Yawson A, Vicente J, Gomes B, Santolamazza F, Moreno M, Charlwood J, Simard F, Elissa N, Weetman D, et al. 2013. Geographic population structure of the African malaria vector Anopheles gambiae suggests a role for the forest-savannah biome transition as a barrier to gene flow. Evol Appl 6:910–924.

Purcell S, Neale B, Todd-Brown K, Thomas L, Ferreira MAR, Bender D, Maller J, Sklar P, de Bakker PIW, Daly MJ, et al. 2007. PLINK: a tool set for whole-genome association and population-based linkage analyses. Am. J. Hum. Genet. 81:559–575.

Robert V, Macintyre K, Keating J, Trape J-F, Duchemin J-B, Warren M, Beier JC. 2003. Malaria transmission in urban sub-Saharan Africa. Am J Trop Med Hyg 68:169–176.

Rougeux C, Bernatchez L, Gagnaire P-A. 2017. Modeling the Multiple Facets of Speciation-with-Gene-Flow toward Inferring the Divergence History of Lake Whitefish Species Pairs (Coregonus clupeaformis). Genome Biology and Evolution 9:2057–2074.

Sabeti PC, Varilly P, Fry B, Lohmueller J, Hostetter E, Cotsapas C, Xie X, Byrne EH, McCarroll SA, Gaudet R, et al. 2007. Genome-wide detection and characterization of positive selection in human populations. Nature 449:913–918.

Sangbakembi-Ngounou C, Ngoagouni C, Akone-Ella R, Kengne P, Costantini C, Nakoune E, Ayala D. 2022. Temporal and biting dynamics of the malaria vectors Anopheles gambiae and Anopheles coluzzii harboring the 2La inversion in Bangui, Central African Republic. Pre-prints Available from: https://www.authorea.com/users/485464/articles/570840-temporal-and-biting-dynamics-of-the-malaria-vectors-anopheles-gambiae-and-anopheles-coluzzii-harboring-the-2la-inversion-in-bangui-central-african-republic?commit=6eea5e83e47b0bedc3788a29130fc0113bb3a695

Schrider DR, Kern AD. 2017. Soft Sweeps Are the Dominant Mode of Adaptation in the Human Genome. Molecular Biology and Evolution 34:1863–1877.

Sharakhova MV, Hammond MP, Lobo NF, Krzywinski J, Unger MF, Hillenmeyer ME, Bruggner RV, Birney E, Collins FH. 2007. Update of the Anopheles gambiaePEST genome assembly. Genome Biology 8:R5.

Sheehan S, Song YS. 2016. Deep Learning for Population Genetic Inference. PLOS Computational Biology 12:e1004845.

Simard F, Ayala D, Kamdem GC, Pombi M, Etouna J, Ose K, Fotsing J-M, Fontenille D, Besansky NJ, Costantini C. 2009. Ecological niche partitioning between Anopheles gambiae molecular forms in Cameroon: the ecological side of speciation. BMC Ecol 9:17.

Slotman MA, Tripet F, Cornel AJ, Meneses CR, Lee Y, Reimer LJ, Thiemann TC, Fondjo E, Fofana A, Traoré SF, et al. 2007. Evidence for subdivision within the M molecular form of Anopheles gambiae. Molecular Ecology 16:639–649.

Small ST, Costantini C, Sagnon N, Guelbeogo MW, Emrich SJ, Kern AD, Fontaine MC, Besansky NJ. 2023. Standing genetic variation and chromosome differences drove rapid ecotype formation in a major malaria mosquito. Proceedings of the National Academy of Sciences 120:e2219835120.

Smith CCR, Tittes S, Ralph PL, Kern AD. 2023. Dispersal inference from population genetic variation using a convolutional neural network. Genetics 224:iyad068.

Smith JM, Haigh J. 1974. The hitch-hiking effect of a favourable gene. Genetics Research 23:23–35.

Tene Fossog B, Antonio-Nkondjio C, Kengne P, Njiokou F, Besansky NJ, Costantini C. 2013. Physiological correlates of ecological divergence along an urbanization gradient: differential tolerance to ammonia among molecular forms of the malaria mosquito Anopheles gambiae. BMC Ecology 13:1.

Tene Fossog B, Ayala D, Acevedo P, Kengne P, Ngomo Abeso Mebuy I, Makanga B, Magnus J, Awono-Ambene P, Njiokou F, Pombi M, et al. 2015. Habitat segregation and ecological character displacement in cryptic African malaria mosquitoes. Evolutionary Applications 8:326–345.

Tennessen JA, Ingham VA, Toé KH, Guelbéogo WM, Sagnon N, Kuzma R, Ranson H, Neafsey DE. 2021. A population genomic unveiling of a new cryptic mosquito taxon within the malaria-transmitting Anopheles gambiae complex. Molecular Ecology 30:775–790.

Tiffin P, Ross-Ibarra J. 2014. Advances and limits of using population genetics to understand local adaptation. Trends in Ecology & Evolution 29:673–680.

Tine M, Kuhl H, Gagnaire P-A, Louro B, Desmarais E, Martins RST, Hecht J, Knaust F, Belkhir K, Klages S, et al. 2014. European sea bass genome and its variation provide insights into adaptation to euryhalinity and speciation. Nat Commun 5:5770.

Tran DT, Zhang L, Zhang Y, Tian E, Earl LA, Ten Hagen KG. 2012. Multiple Members of the UDP-GalNAc: Polypeptide N-Acetylgalactosaminyltransferase Family Are Essential for Viability in Drosophila. J Biol Chem 287:5243–5252.

Venkatesan P. 2024. The 2023 WHO World malaria report. The Lancet Microbe 5:e214.

Verdu P, Austerlitz F, Estoup A, Vitalis R, Georges M, Théry S, Froment A, Le Bomin S, Gessain A, Hombert J-M, et al. 2009. Origins and Genetic Diversity of Pygmy Hunter-Gatherers from Western Central Africa. Current Biology 19:312–318.

Visser ME. 2008. Keeping up with a warming world; assessing the rate of adaptation to climate change. Proceedings of the Royal Society B: Biological Sciences 275:649–659.

Voight BF, Kudaravalli S, Wen X, Pritchard JK. 2006. A Map of Recent Positive Selection in the Human Genome. PLOS Biology 4:e72.

White BJ, Collins FH, Besansky NJ. 2011. Evolution of Anopheles gambiae in Relation to Humans and Malaria. Annual Review of Ecology, Evolution, and Systematics 42:111–132.

Willis KJ, Bennett KD, Burrough SL, Macias-Fauria M, Tovar C. 2013. Determining the response of African biota to climate change: using the past to model the future. Philosophical Transactions of the Royal Society B: Biological Sciences 368:20120491.

Zohdy S, Derfus K, Headrick EG, Andrianjafy MT, Wright PC, Gillespie TR. 2016. Small-scale land-use variability affects Anopheles spp. distribution and concomitant Plasmodium infection in humans and mosquito vectors in southeastern Madagascar. Malaria Journal 15:114.

