## Supplementary Figures for "Genomic Signatures of Microgeographic Adaptation in *Anopheles coluzzii* Along an Anthropogenic Gradient in Gabon"

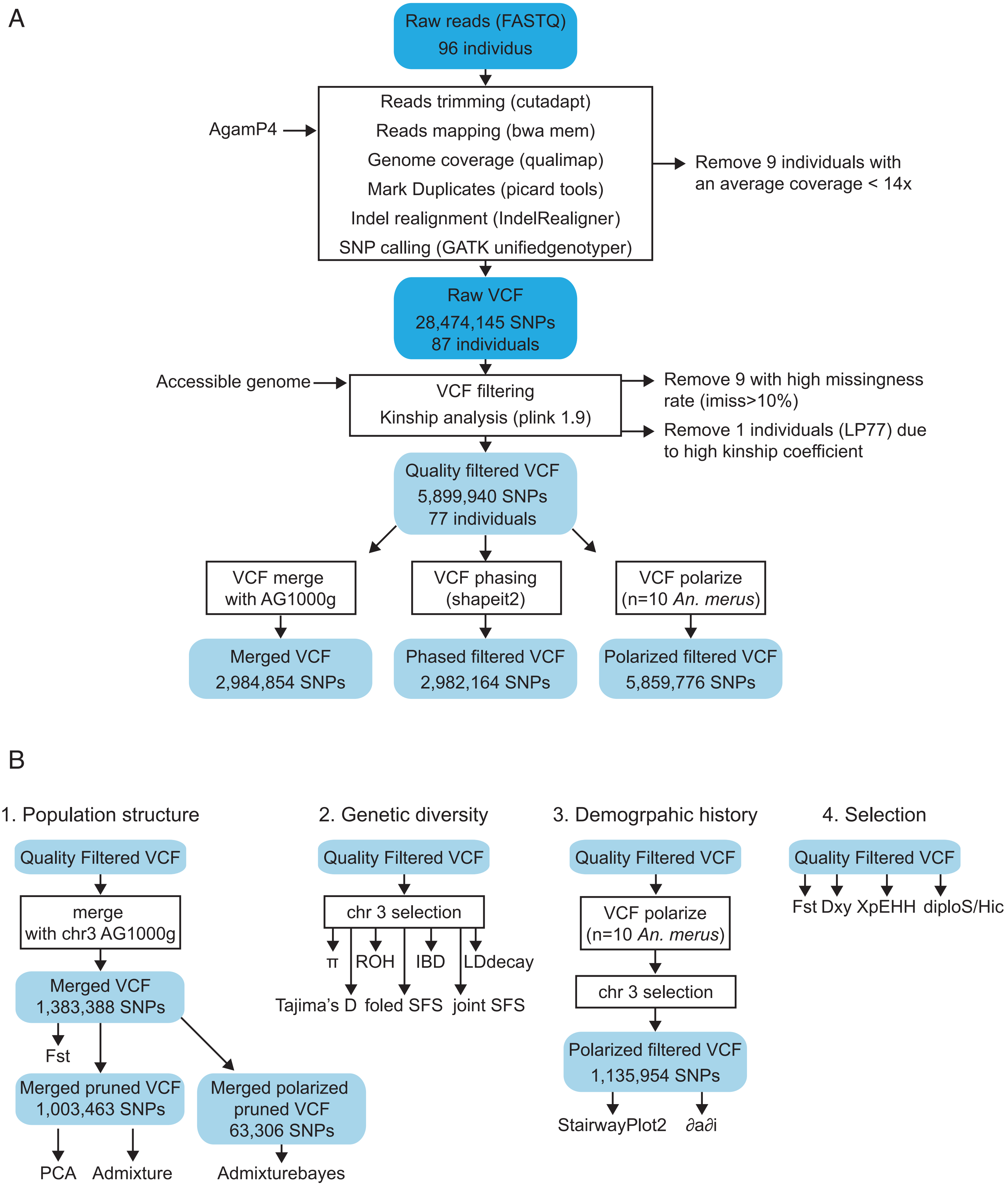


**Supplementary Figure 1:** (A) Workflow of the reads mapping and SNP genotyping procedures used. (B) Schematic overview of the different analysis and the input dataset used.

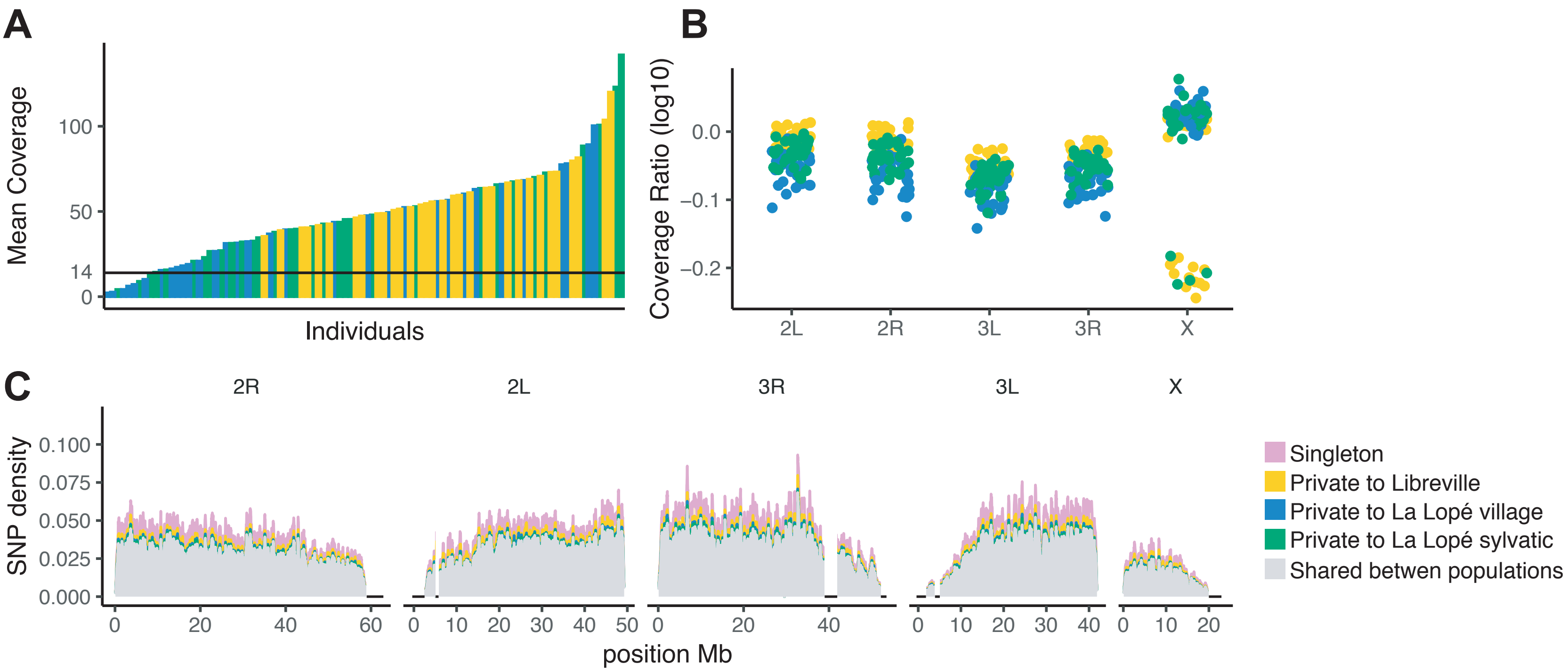


**Supplementary Figure 2: Sequencing coverage depth and SNP density along the genome.** (**A**) Distribution of the mean sequencing depth of the 96 samples. Bars represent individual mosquito samples and are color-coded according to their sampling origin: Libreville (yellow); La Lopé village (blue); and La Lopé sylvatic (green). The horizontal line represents the coverage cut-off at 14x used to exclude samples below that threshold. (**B**) Coverage ratio of the mean sequencing depth for each chromosome over the coverage of the whole genome. This allowed to assign the sex of each individual, which was unknown for the larvae from Libreville (see Supplementary Table S1). (**C**) Density of the high-quality SNPs in 200-kb non-overlapping windows over the genome.

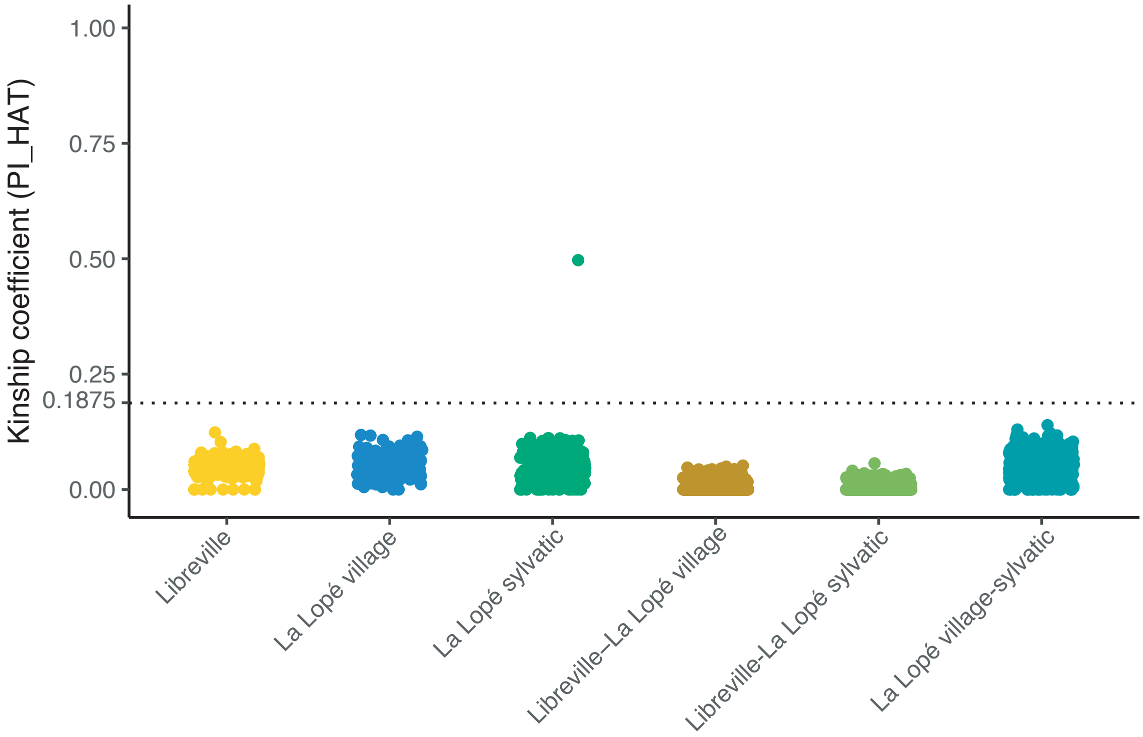


**Supplementary Figure 3: Kinship analysis across the Gabonese dataset estimated with pair-wise IBD estimator (PI_HAT) between samples in PLINK.** The threshold 0.1875 represents the half-way point between 2nd and 3rd degree relatives and is a common cut-off to use.

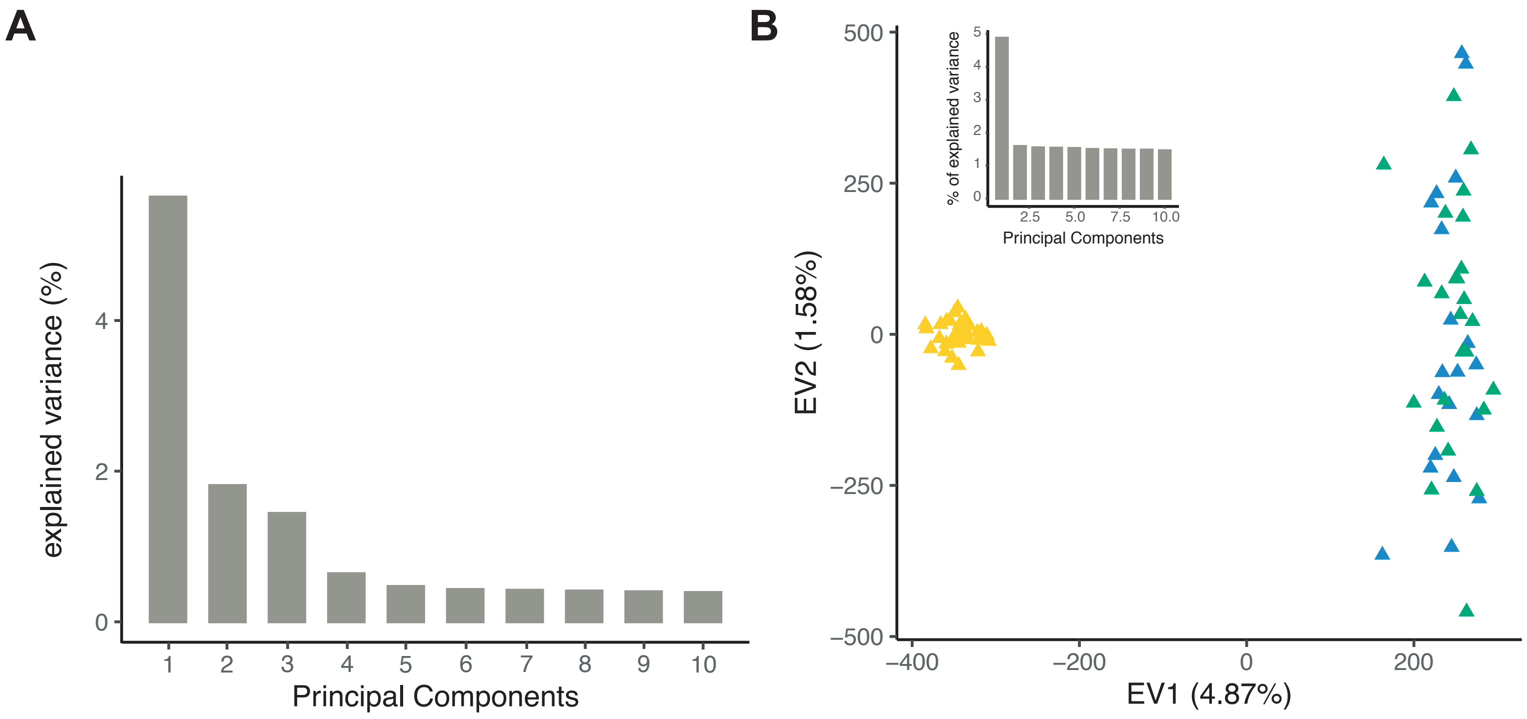


**Supplementary Figure 4:** (**A**) Scree-plot showing the variance fraction explained by each principal component of the PCA for the African *An. coluzzii* samples (combining the Gabonese and AG1000G datasets) represented in Figure 1. (**B**) PCA of the 77 *An. coluzzii* mosquitoes from Gabon retained for further analysis using biallelic SNPs from the euchromatic regions of the chromosome 3. The bar chart shows the percentage of variance explained by each principal component.

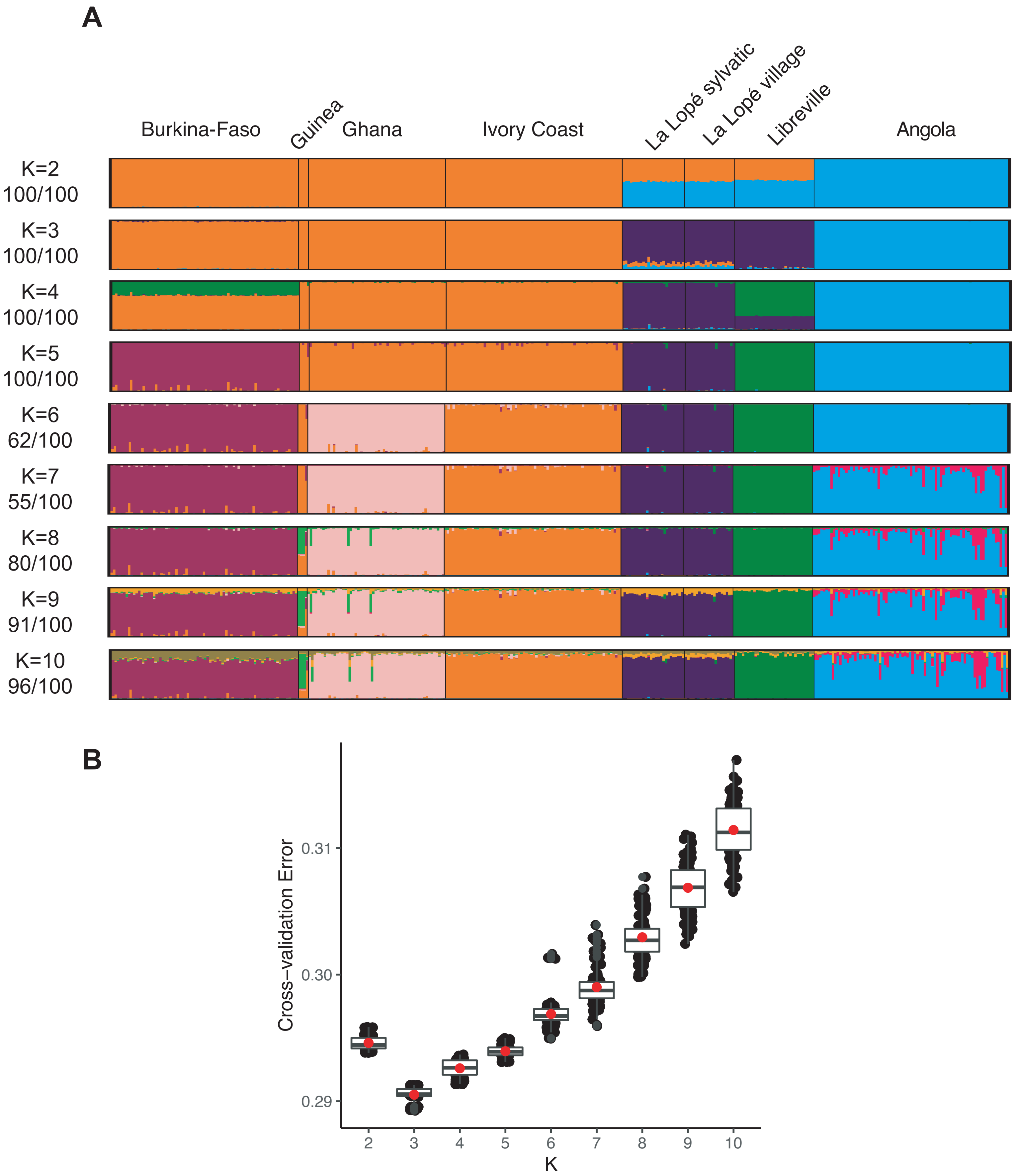


**Supplementary Figure 5: Analysis of population structure and genetic ancestry in *An. coluzzii* considering the Gabonese populations in perspective with those from the Ag1000G.** (**A**) Individual ancestry proportions (from K=2 to K=10) were estimated using the ADMIXTURE program. Each vertical bar represents an individual mosquito grouped according to sampling location and colored according to the proportion of the genome inherited from each of the K ancestral clusters tested. (**B**) Box-plots showing the average (red dots), median, and interquartile values of the cross-validation (CV) error rate estimated using the ADMIXTURE program for each ancestral cluster tested (with K ranging between 2 and 10). Black dots show the CV error rate values for 100 replicated runs at each tested K values. K=3 was chosen as the best-fitted solution for our SNP dataset, since that value minimizes the CV error rate.

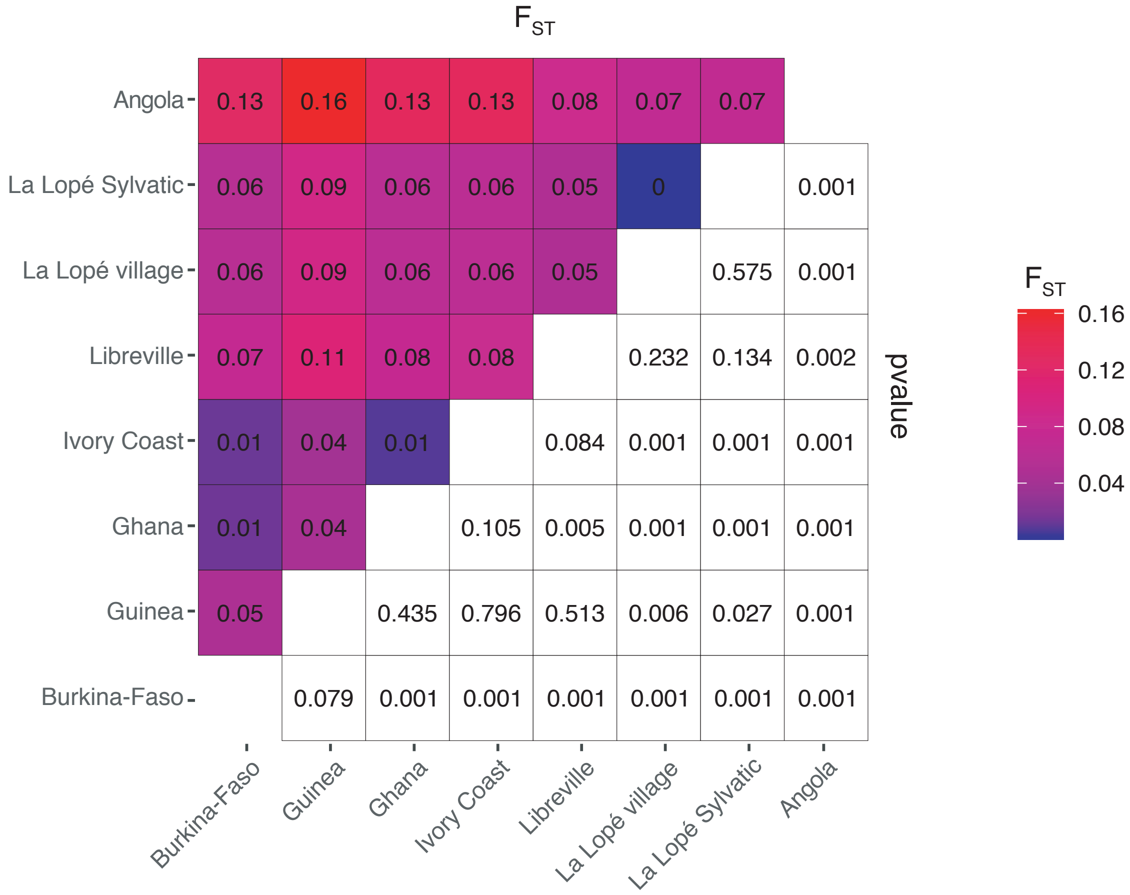


**Supplementary Figure 6: Pairwise population differentiations ($F_{ST}$) among populations of *An. coluzzii*.** Average differentiation in allele frequency estimated using the $F_{ST}$ statistics between pairs of populations. Upper left portion of the matrix shows average $F_{ST}$ values between each population pair. Bottom right portion of the matrix shows the p-value derived from the z-score for each $F_{ST}$ value estimated via a block-jackknife procedure.

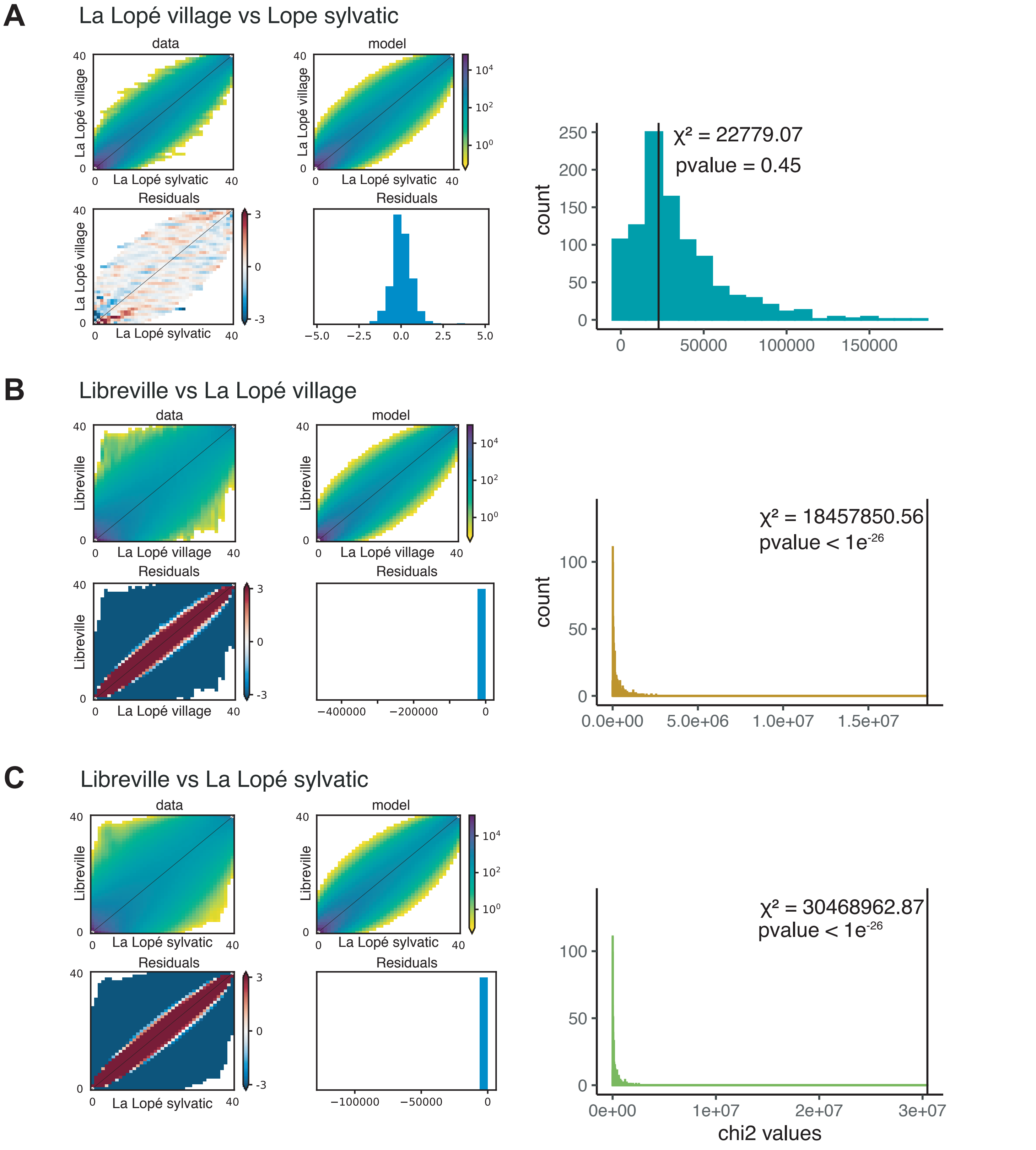


**Supplementary Figure 7: Test of departure from random mating expectation (panmixia) between pairs of populations performed using δaδi.** The left panels represent the observed joint site frequency spectrum (jSFS) between populations pairs along with a model fit and residuals using δaδi, for a “scramble” model where individuals are permuted across population. On the right panel, the null distribution of χ2 values is obtained by measuring the deviation between 1000 replicates of permuted individual labels across the population pair to the scramble model. Vertical black line and value correspond to the χ2 value calculated between the observed jSFS and the scramble jSFSmodel. This test of departure from panmictic expectation was performed for all population pairs including : La Lope village versus La Lope sylvatic (**A**), Libreville versus La Lope village (**B**), and Libreville versus La Lope sylvatic (**C**).

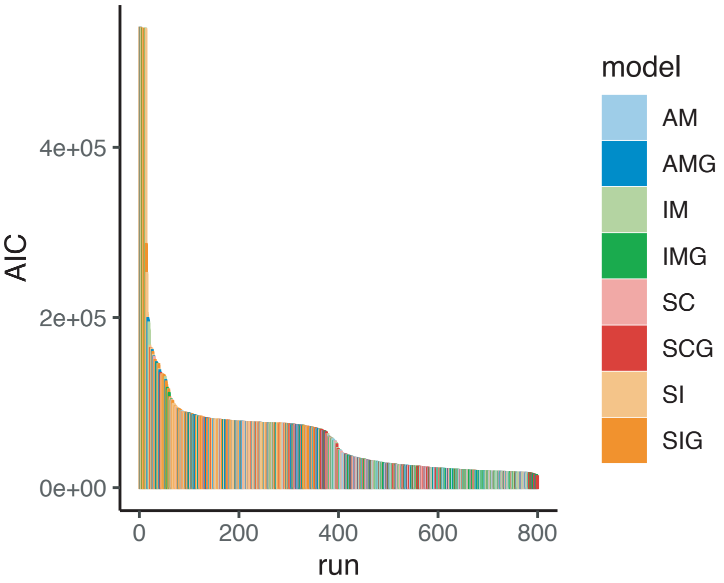


**Supplementary Figure 8: δaδi model selection based on the AIC score obtained for 8 different models with 100 replicates.** The lowest AIC score was observed for the model secondary contact with growth (SCG).

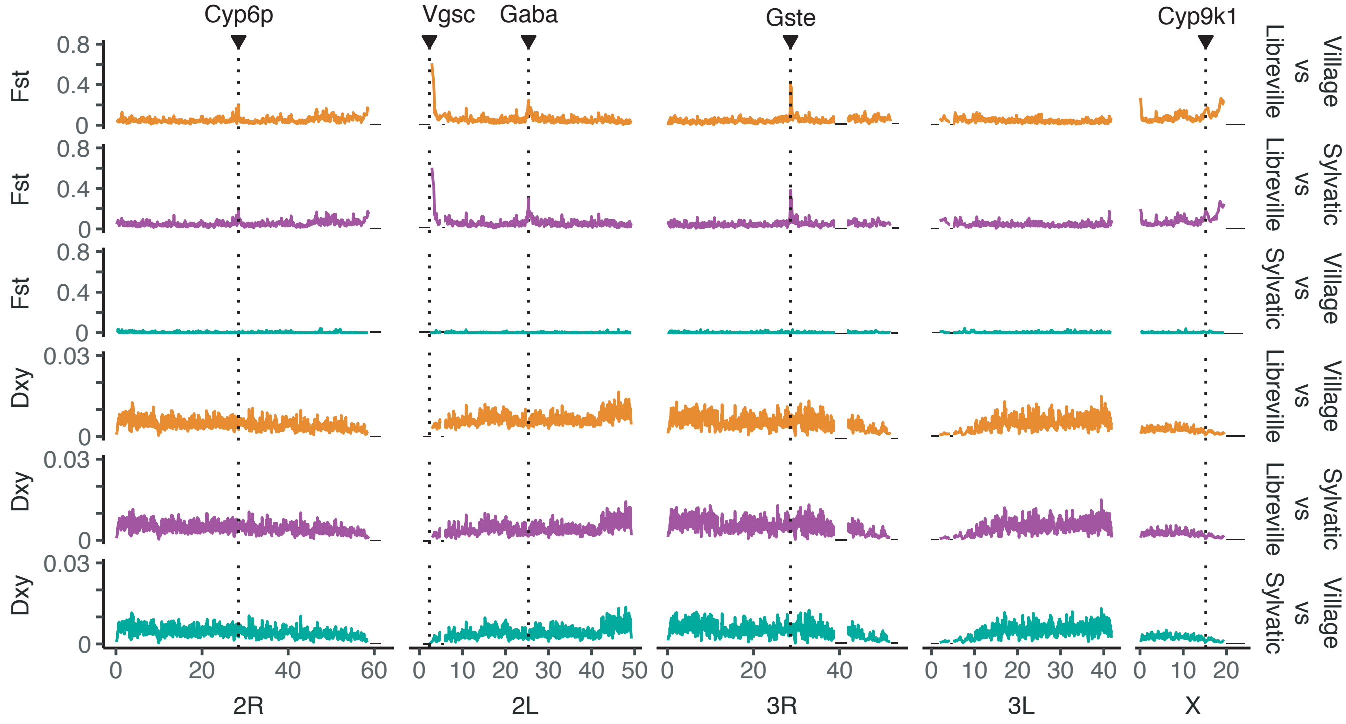


**Supplementary Figure 9: Genome scan of $F_{ST}$ and $D_{xy}$ statistics.** Both statistics were calculated in 1000kb non-overlapping windows and plotted for all pairwise comparisons. Fine horizontal black lines indicate the heterochromatic regions excluded from the analyses, and dotted lines indicate known insecticide resistance genes.

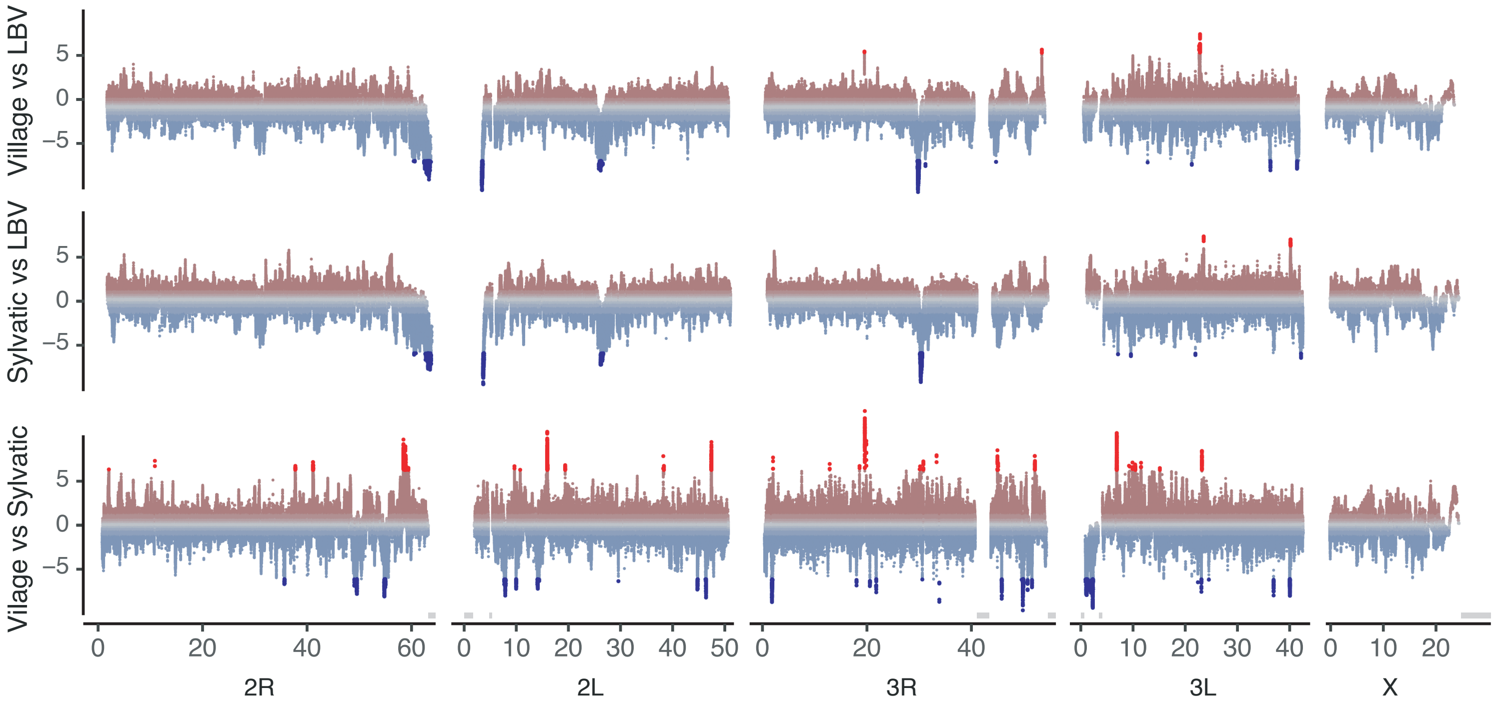


**Supplementary Figure 10: Genome scan of XP-EHH scores calculated at the SNP level along the genome and plotted for all pairwise population comparisons.** For each population comparison (e.g., LLP village vs LBV), positive scores indicate longer haplotype homozygosity and therefore recent selection in the first population (e.g., LLP village), and negative scores indicate selection in the second population (e.g., LBV). Each dot has been colored by its associated p-value for the XP-EHH score, with shaded red and blue colors gradient representing non-significant SNPs, and red and blue representing significant SNPs (p-value < 1e-4). Gray areas above the x-axis indicate heterochromatic regions excluded from our analysis, and dotted lines indicate candidate genes including known insecticide genes.

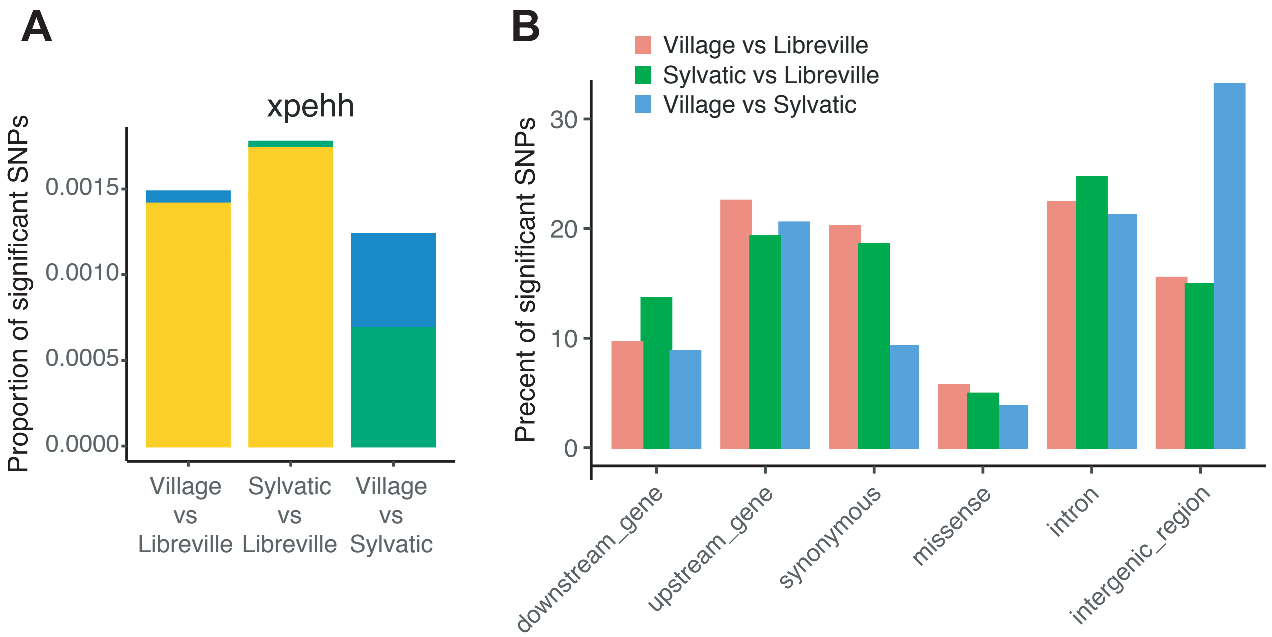


**Supplementary Figure 11:** (**A**) Barplot representing the proportion of SNPs displaying significant p-value for the XP-EHH score for each of the 3 pairwise population comparisons. Color code represents the population in which the SNP has been found significant (Yellow – LBV; Green – LLP sylvatic; Blue – LLP village). (**B**) Functional annotation of the SNPs identified as significant using the XP-EHH scores in each of the three pairwise comparisons.

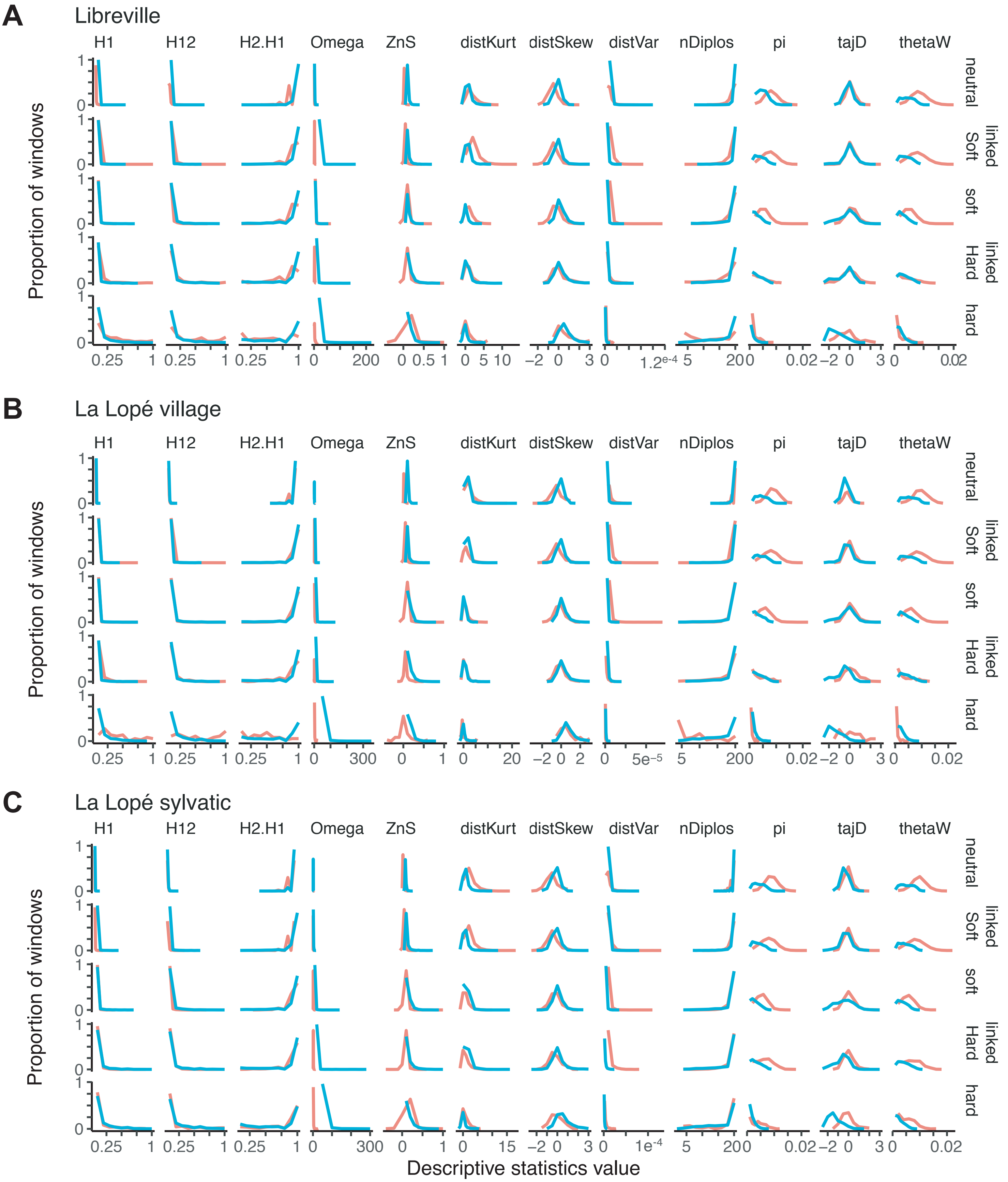


**Supplementary Figure 12: Goodness-of-fit between empirical and simulated data under the 5 different types of selection scenarios of selective sweep in the *diploS/HIC* analysis.** The distributions obtained for each of the 12 summary statistics used in *diploS/HIC* and for each population (LBV, LPV, LPS) are displayed for the simulated (red) and empirical (bleu) data.

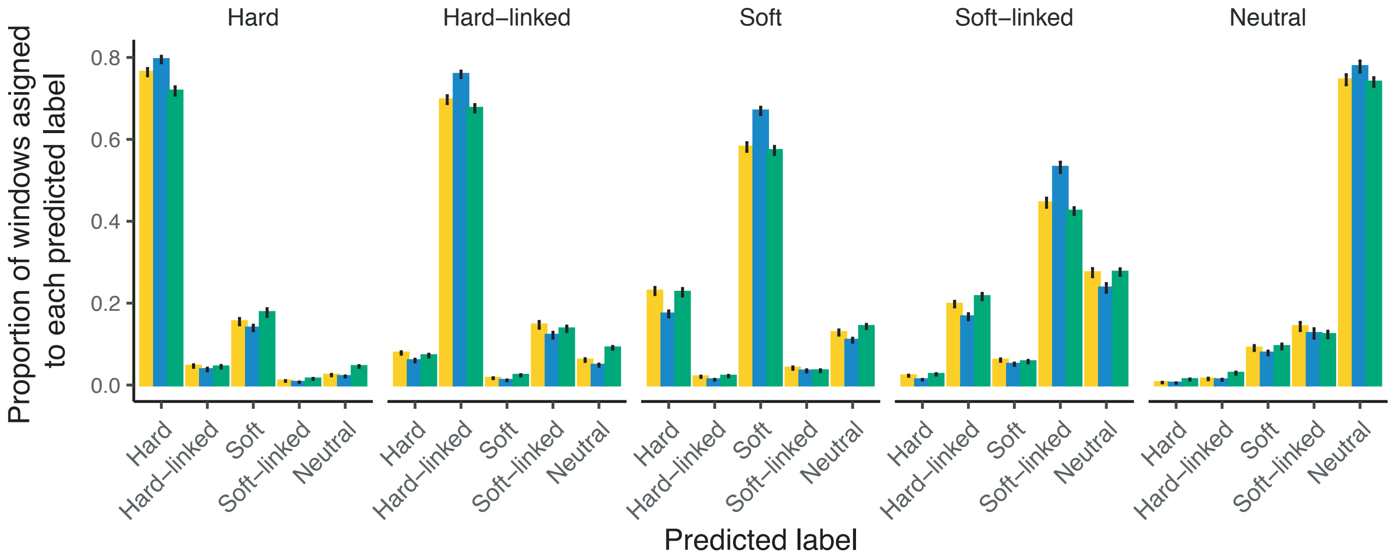


**Supplementary Figure 13: Graphical representation of the confusion matrix.** Each matrix is represented in form of a barplot, with each facet of the figure represent the true label of each testing set, the x-axis represents the predicted type of selective sweep and the y-axis the proportion of windows sign to each sweep type. Each barplot consist of the average of the 100 replicates with the confidence interval represented as error bar.
